# Kremen1 regulates the regenerative capacity of support cells and mechanosensory hair cells in the zebrafish lateral line

**DOI:** 10.1101/2023.07.27.550825

**Authors:** Ellen Megerson, Michael Kuehn, Ben Leifer, Jon Bell, Hillary F. McGraw

**Affiliations:** Division of Biological and Biomedical Systems, School of Science and Engineering, University of Missouri-Kansas City, Kansas City, MO 64110, USA; Department of Cancer Biology, University of Kansas Medical Center, Kansas City, KS 66103, USA; Department of Population Health, University of Kansas Medical Center, Kansas City, KS 66103, USA

## Abstract

Mechanosensory hair cells in the inner ear mediate the sensations of hearing and balance, and in a specialize lateral line sensory system of aquatic vertebrates, the sensation of water movement. In mammals, hair cells lack the ability of regenerate following damage, resulting in sensory deficits. In contrast, non-mammalian vertebrates, such zebrafish, can renew hair cells throughout the life of the animal. Wnt signaling is required for development of inner ear and lateral line hair cells and regulates regeneration. Kremen1 inhibits Wnt signaling and hair cell formation, though its role in regeneration has not been established. We use a zebrafish *kremen1* mutant line, to show that when Wnt signaling is overactivated in the lateral line, excessive regeneration occurs in the absence of increased proliferation, due to an increase in support cells. This contrasts with the previously described role of Wnt signaling during hair cell regeneration. This work will allow us to understand the biology of mechanosensory hair cells, and how regeneration might be promoted following damage.

## Introduction

Loss of hearing is one the most common sensory defects found among the human population; an estimated 15% of the world’s population has some degree of hearing loss and over 5% have disabling loss of auditory function (https://www.who.int/news-room/fact-sheets/detail/deafness-and-hearing-loss). Hearing and vestibular functions are mediated by specialized mechanosensory hair cells found within the inner ear. In mammals, when the hair cells are damaged through exposure to noise, ototoxic drugs, injury, or disease, they fail to regrow (^1–5^). In contrast, non-mammalian vertebrates, such as chickens, frogs, and fish, can regenerate hair cells, in cases throughout their lifespan (^6–8^). Understanding how hair cells regenerate in some animals is critical to identifying potential therapies for human hearing loss.

In addition to inner ear hair cells, aquatic vertebrates have mechanosensory hair cells in their lateral lines systems, which mediate sensations of water movement (^9^). Lateral line hair cells are morphologically and genetically very similar to the inner ear hair cells of the auditory and vestibular systems (^10^). Unlike the inner ear, which is difficult to access and experimentally manipulate, the lateral line is arrayed on the surface of the body and amenably to manipulation (^11, 12^). The zebrafish has emerged as an excellent model organism for the study of mechanosensory hair cell development and regeneration (^13^). The zebrafish lateral line comprises sensory organs, called neuromasts, which contain hair cells, which sense water movement, and surrounding support cells, which act as stem cells to maintain neuromast homeostasis (^14^). In the zebrafish, lateral line hair cells are robustly regenerative throughout the life of the animal (^6^).

Recent work has demonstrated that within the cohort of neuromast support cells there are specific populations which can preferentially give rise to regenerated hair cells, self-renew to maintain stem cell numbers, or remain quiescent under most conditions (^15–19^). Single-cell RNA-sequencing has identified specific gene expression profiles for the distinct populations of neuromast support cells during hair cell regeneration (^15, 16, 18, 20^). These transcriptional profiles served as the basis for cell labeling experiments to analyze the behavior of support cells as hair cells are regrown. In particular, the *Tg(sost:nlsEos)^w216^* transgenic line allowed conditional labeling of dorsoventral support cells and demonstrated that during regeneration these cells proliferate and give rise to the majority of hair cells (^19^). The proliferation of anterior-posterior support cells labeled with the *Tg(tnfs10l3:nlsEos)^w223^* formed the minority of regenerating hair cells, and primarily gave rise to self-renewing support cells. Finally, peripheral support cells labeled with *Tg(sfrp1a:nslEos)^w217^* were largely quiescent, though studies have shown that extensive ablation of interior support cells can trigger peripheral support cells to proliferate and repopulate the neuromast (^19, 21, 22^). As we gain a better understanding of how specific populations of support cells contribute to regeneration, it raises the question of how these cells are specified?

Fgf, Notch and Wnt signaling, have been found to regulate proliferation and regeneration in the zebrafish lateral line (^16, 23–26^). The Fgf and Notch pathways regulate the maintenance of proper hair cell number and have been found to inhibit support cell proliferation and hair cell differentiation under homeostatic conditions. Blocking the function of either Fgf or Notch during neuromast regeneration following hair cell ablation, either using pharmacological antagonists or in mutant zebrafish lines, results in increased support cell proliferation and the differentiation of supernumerary hair cells ^16, 19, 24^). Analysis of cell sub-populations found that inhibiting Notch signaling results in an increase in regenerated hair cells forming from dorsoventral and anterior-posterior populations. In particular, *Tg(tnfs10l3:nlsEos)^w223^* contributed to a greater proportion of regenerated hair cells as compared to control regeneration conditions (^19^).

Canonical Wnt signaling has been implicated as a critical pathway regulating hair cell regeneration in the zebrafish lateral line. Studies examining gene expression patterns and single cell RNA-sequencing during neuromast regeneration revealed that canonical Wnt signaling is upregulated beginning 3 hours after hair cell ablation and remains active during the first 10 hours of regeneration (^15, 16, 24^). Studies using pharmacological manipulation of the Wnt pathway or overexpression of pathway inhibitors such as heat-shock induction of Dkk1b, suggest that Wnt signaling is required for the proliferation of support cells and hair cell regeneration (^16, 24, 27^). In these studies, blocking the Wnt pathway results in the regeneration of fewer hair cells, while activating Wnt signaling results in increased proliferation and hair cell regeneration. Our current study will be the first to specifically examine regeneration in a Wnt pathway mutant.

Kremen1 (Krm1) is a vertebrate-specific member of the Wnt pathway which inhibits Wnt signaling as a co-receptor for the secreted family of Dkk proteins (^28, 29^). In the mouse cochlea, *krm1* is expressed in support cells and inhibiting Krm1 function in cochlear explants results in the formation of excess and disordered hair cells, but not significant changes in proliferation (^30^). In zebrafish, a *krm1* mutant line (*krm1^nl10^*) shows truncated posterior lateral line formation and an increase in the development of neuromast hair cells (^30, 31^). Blocking *krm1* function with a morpholino, resulted in the development of physically larger neuromasts, though this study did not find a significant change in hair cell number (^32^). It has not been established if Krm1 function plays a role in hair cell regeneration.

In this study we will use the *krm1^nl10^* zebrafish mutant line to determine the role of Krm1 in the regulation of cellular behavior during regeneration in the zebrafish posterior lateral line (^31^). Our analysis of hair cell numbers following ablation and regeneration shows a significant increase in *krm1^nl10^* mutants, though not an increase in cellular proliferation. Instead we find that loss of Krm1 function results in an increase of *Tg(sost:nlsEos)^w216^* labeled dorsoventral support cells, which give rise to regenerated hair cells. We find that we can eliminate the supernumerary hair cell in *krm1^nl10^* larvae following repeated bouts of regeneration, suggesting that there is a poised population of support cells which can be overwhelmed by repeat regeneration. Finally, we demonstrate that inhibition of Notch signaling further bias the number of regenerated hair cells in *krm1^nl10^* mutant larvae as well as increases in proliferation and in dorsoventral support cells, suggesting that the Notch and Wnt signaling may act independently to regulate cellular differentiation and proliferation. These results suggest that regulation of Krm1 function is required for proper hair cell renewal after damage.

## Results

### *krm1* is dynamically expressed during regeneration in posterior lateral line NMs

Previous research demonstrated that *krm1* is expressed in migrating posterior lateral line primordium during the initial 48 hours of development in the zebrafish embryo and in the support cells of developing adult mouse cochlea (^30, 31^). To determine the expression patterns on *krm1* in the larval zebrafish lateral line during regeneration, we used whole mount RNA in situ hybridization (WISH ^33^) and hybridization chain reaction (HCR^34, 35^) for fluorescent in situ hybridization (FISH). At 5 dpf, prior to NEO-induced hair cell ablation, *krm1* is expressed throughout the neuromast (Fig. 1A), when Wnt signaling is expected to be low (^24^). To determine the expression pattern of *krm1* during hair cell regeneration, we examined 4 hours post NEO neuromasts (Fig. 1B), when Wnt is upregulated (^15, 16, 24^) and 1 day post NEO (Fig. 1C), at these time points *krm1* expression is greatly reduced. By 3 days post NEO, regeneration is complete and *krm1* expression is upregulated in neuromasts (Fig. 1D). Building on the dynamic expression of *krm1* during regeneration, we next sought to determine if there is cell-specific localization of *krm1* in NMs. To do this we used HCR-FISH in *Tg(myosin6b:GFP)^w186^* (*myo6:GFP*) expressing larvae. We found that at 5dpf HCR *krm1* is expressed throughout the neuromast in surrounding support cells and in *myo6:GFP*-positive hair cells (E-E’’’). In agreement with our WISH experiments, at 4 hours post NEO-exposure we see an absence of HCR *krm1* expression (F-F’’’) and a moderate level of expression by 1-day post NEO (G-G’’’). By 3-days after NEO-exposure, HCR *krm1* expression is up regulated in regenerated NMs (H-H’’’). These results indicate that *krm1* expression is present during homeostasis in NMs, when Wnt signaling is low and downregulated during regeneration when Wht signaling is upregulated.

**Figure 1.**
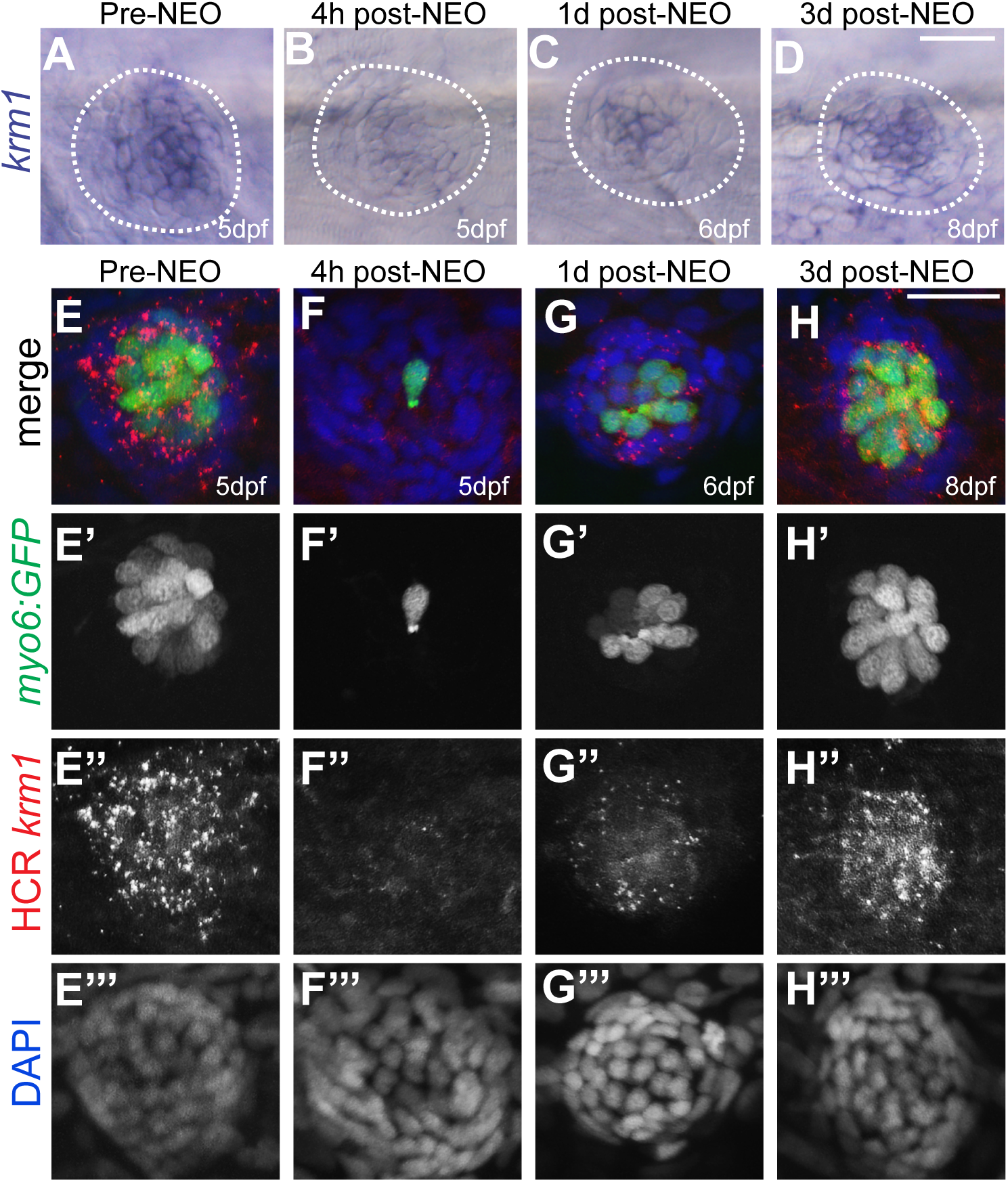
*krm1* expression is dynamic during posterior lateral line regeneration. (**A-D**) RNA in situ hybridization of *krm1* showing expression in 5dpf NM prior to exposure to NEO, 4h after NEO exposure,1-day post NEO exposure in a 6dpf larva, and 3-days post NEO exposure. Scale bar=20μm. (**E-H**) Confocal projections of HCR-FISH showing *krm1* expression (red) during regeneration in wild-type NMs in *Tg(myosin6b:GFP)^w186^* larvae (green) with nuclei labeled with DAPI (blue) in 5dpf NM prior to exposure to NEO, 4h after NEO exposure, 1-day post NEO exposure in a 6dpf larva, and 3-days post NEO exposure. Scale bar=20μm.

### Supernumerary hair cells form during regeneration in *krm1^nl10^* mutants

During development, Krm1 is expressed in the support cells of the mouse cochlea (^30^). When Krm1 function is blocked by RNAi, there is a significant increase in the development of cochlear hair cells, though without a change in levels of proliferation (^30^). Examination of the *krm1^nl10^* zebrafish mutant line also showed an increase in the development of lateral line hair cells (^30^). We sought to determine if Krm1 activity also regulates the regeneration of hair cells in the zebrafish posterior lateral line. At 5dpf, prior to hair cell ablation with NEO, there are significantly more hair cells labeled with *myo6:GFP krm1^nl10^* mutant larvae as compared to heterozygous and wild-type siblings (Fig.2 A,A’,B,B’,G). Total cell numbers as labeled by DAPI and α-Sox2 antibody-labeled support cells within NMs did not show a significant change between mutants and heterozygous sibling controls (A’’-C’’’,H,I). To confirm that hair cells are susceptible to NEO damage, we examined control and *krm1^nl10^* NMs 1-hour post exposure to NEO and found the majority of hair cells were ablated (C-D’’’, G). To assess regeneration, we determined hair cell numbers 3-days post NEO exposure and found a significant increase in *myo6:GFP+* cells in *krm1^nl10^* mutants as compared to heterozygous siblings (E-F’, G), though support cells and total cell numbers were not significantly different (E’’-F’’’, H,I). We also compared the index of hair cell numbers to total cells within NMs to confirm that the differences we are seeing are not the result of changes in overall neuromast size, and found that we still find a significant difference in homeostatic and regenerated hair cells in *krm1^n^*^10^ mutants as compared to heterozygous siblings (J).These results suggest that mechanisms that lead to excess hair development in *krm1^nl10^* mutants may also regulate hair cell regeneration.

**Figure 2.**
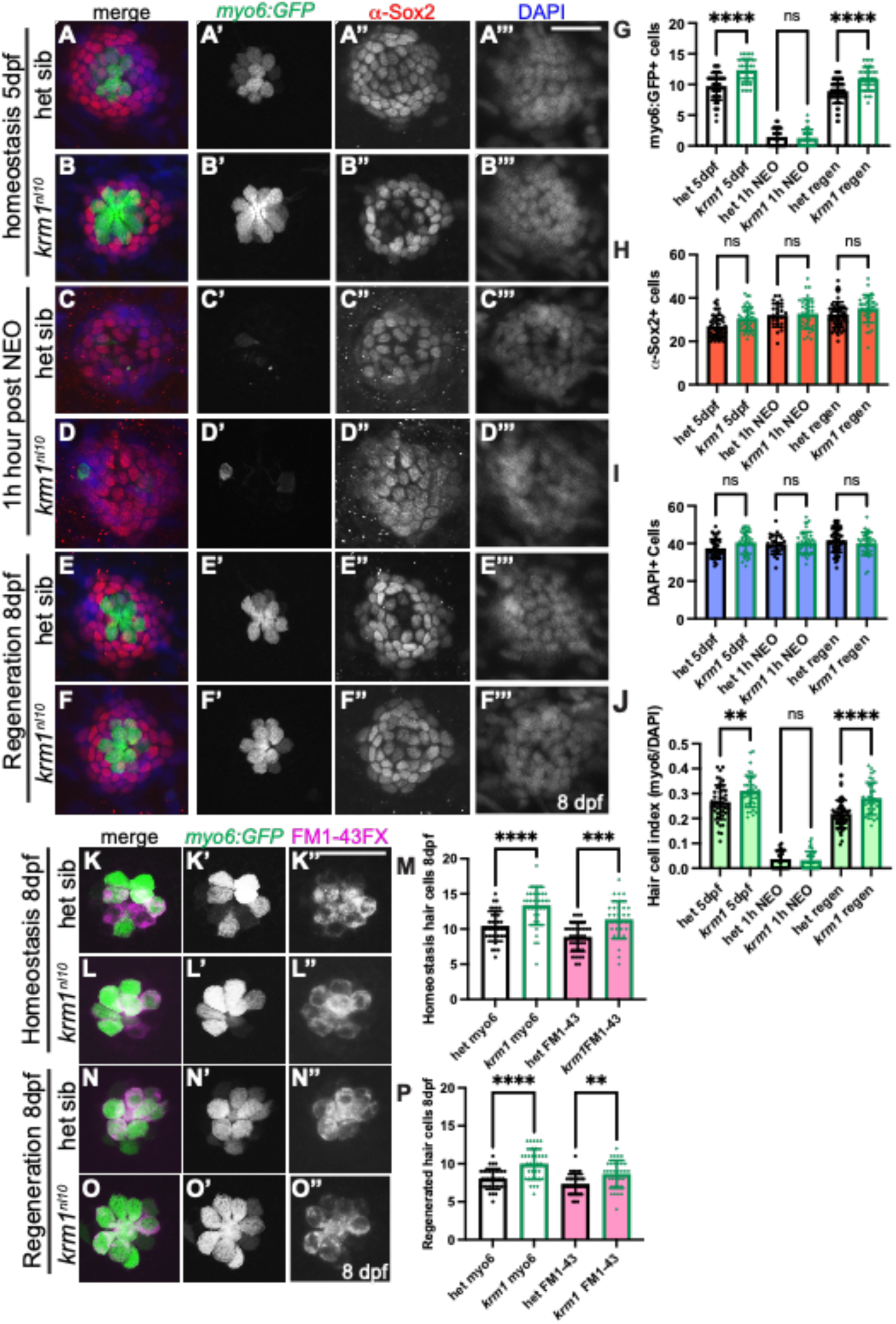
*krm1^nl^*^10^ larvae form supernumary hair cells following regeneration in posterior lateral line neuromasts. (**A-B’’’**) confocal projections of neuromasts in heterozygous sibling and *krm1^nl10^* larvae with *Tg(myosin6b:GFP)^w186^* expression in hair cells (green), α-Sox2 antibody-labeled support cells (red) and DAPI-labeled nuclei (blue) in heterozygous sibling and *krm1^nl10^* larvae with *Tg(myosin6b:GFP)^w186^* expression in hair cells (green), α-Sox2 antibody-labeled support cells (red) and DAPI-labeled nuclei (blue) in heterozygous sibling and *krm1^nl10^* mutant at 5dpf NM prior to exposure to NEO. (**C-D’’’**) Heterozygous sibling and *krm1^nl10^* mutant at 1-hour post NEO (**E-F’’’**) heterozygous sibling and *krm1^nl10^* mutant 3-days post NEO exposure. (**G-J**) Quantification of *Tg(myosin6b:GFP)^w186^* positive hair cells at 5dpf pre-NEO exposure, 1-hour post NEO exposure, and at 8dpf, 3-days post NEO exposure. (**H**). Quantification of α-Sox2 antibody-labeled support cells at 5dpf pre-NEO exposure, 1-hour post NEO exposure, and at 8dpf, 3-days post NEO exposure, DAPI-labeled nuclei at 5dpf pre-NEO exposure, 1-hour post NEO exposure, and at 8dpf, 3-days post NEO exposure, and an index of hair cells *Tg(myosin6b:GFP)^w186^* -positive cells/DAPI-labeled cells) at 5dpf pre-NEO exposure, 1-hour post NEO exposure, and at 8dpf, 3-days post NEO exposure.). 5dpf: het sibling n=48 NMs (9 fish), *krm1^nl10^* n=38 NM (9 fish); 1h-post NEO: het sibling n=25 NMs (7 fish), *krm1^nl10^* n=36 NM (7 fish); 8dpf: het sibling n=57 NMs (11 fish), *krm1^nl10^* n=34 NM (10 fish). (**K-L’’**) Confocal projections of live images showing FM1-43 incorporation (magenta) in *Tg(myosin6b:GFP)^w186^*–positive hair cells (green) at 8dfp in heterozygous sibling *krm1^nl10^* mutant NMs under homeostatic conditions. (**M**) Quantification of labeled hair cells during homeostasis, heterozygous sibling n=36 NM (9 fish), *krm1^nl10^* n=31 NM (10 fish). (**N-O’’**) FM1-43 incorporation (magenta) in *Tg(myosin6b:GFP)^w186^*–positive hair cells (green) at 8dfp heterozygous sibling and *krm1^nl10^* mutant NMs under regeneration conditions. (**P**) Quantification of regenerated hair cells, heterozygous sibling n=34 NM (10 fish), *krm1^nl10^* n=33 NM (10 fish). All date presented as mean ±SD, \*\**p*=0.006, \*\*\**p*=0.0003, \*\*\*\**p*<0.0001 one-way ANOVA, Tukey’s multiple comparisons test. Scale bar=20μm.

We next sought to confirm that the supernumerary hair cells found in *krm1^nl10^* mutants are functional. Do to this, we used the vital dye FM1-43FX (Thermo Fisher) which enters the cell through functional mechanotransduction channels (^36, 37^). Significantly more hair cells are present at 8dpf in *krm1^nl10^* mutant NMs under both homeostatic and regeneration conditions (K-K’’, M-M’’,N,O) as compared to heterozygous siblings (J-J’’, L-L’’, N,O). Together these results suggest that loss of Krm1 function results in the formation of supernumerary functional posterior lateral line hair cells during development and regeneration.

### Wnt activity is responsible for excess hair cells *krm1^nl10^* mutant NMs

The canonical Wnt signaling pathway is activate during hair cell regeneration in the zebrafish lateral line (^24^). Single-cell RNA sequencing studies have demonstrated that Wnt signaling is upregulated beginning 3 hours after hair cell ablation with NEO in the zebrafish lateral line (^15, 16^). To assess Wnt signaling dynamics during regeneration in our *krm1^nl10^* line, we examined the expression of *β-catenin b1* (*ctnnb1*) and *wnt2* by RNA in situ hybridization in 5 dpf larvae prior to NEO-exposure and 4-hours post NEO-exposure. We found low levels of *ctnnb1* and *wnt2* expression in wild-type and mutant NMs at 5dpf (Fig. 3A-B,E-F) and increased expression of both RNAs at 4-hours post NEO in wild-type and *krm1^nl10^* mutant larvae (Fig. 3 C-D,G-H). These results indicate that Wnt signaling is active during regeneration in control and *krm1^nl10^* NMs.

**Figure 3.**
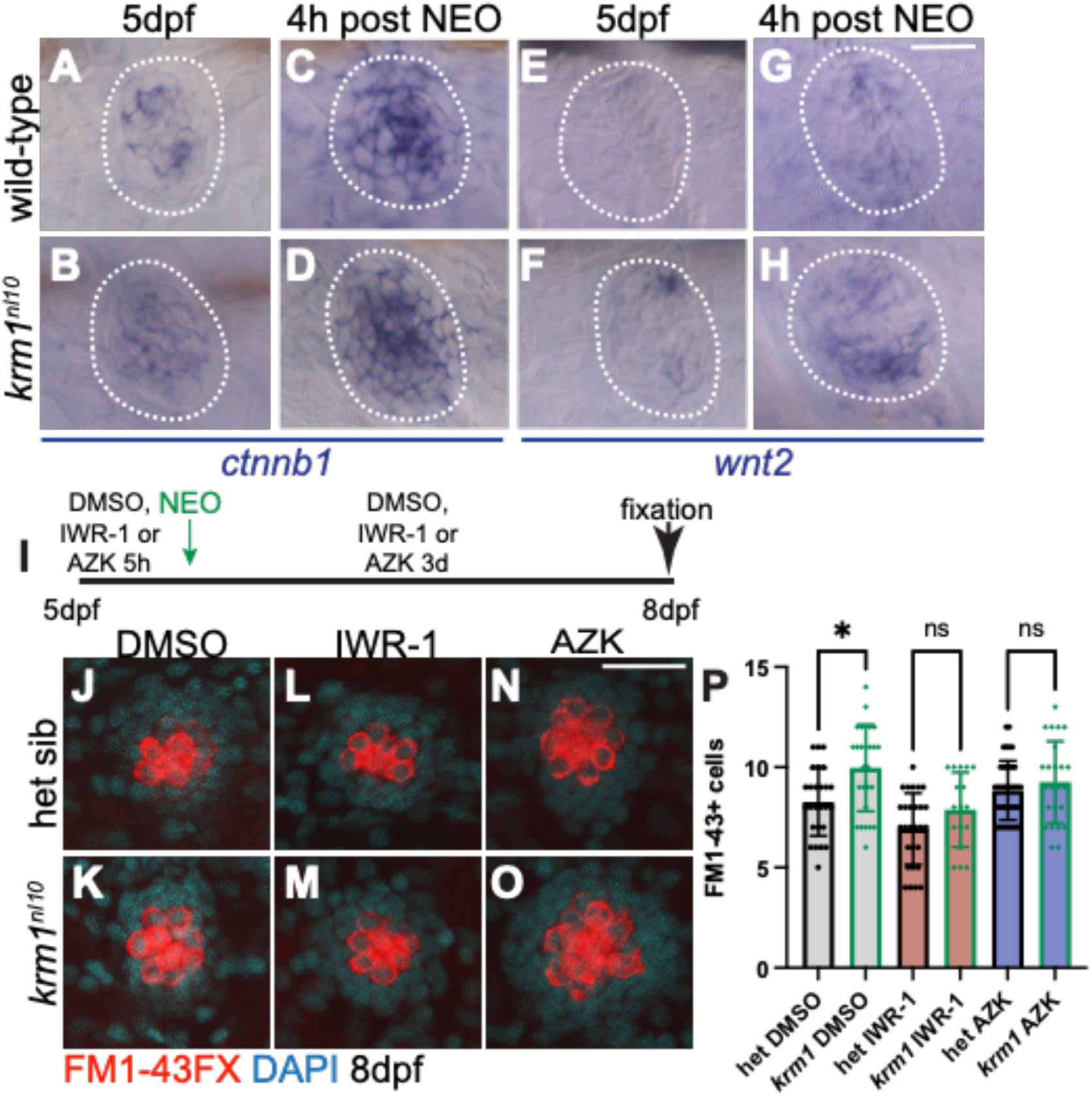
Wnt signaling is active during regeneration in *krm1^nl^*^10^ neuromasts. (**A-D**) Representative images of RNA in situ hybridization in at 5dpf of *ctnnb1* expression in wild-type and *krm1^nl10^* mutant NMs without exposure to NEO and 4 hours post NEO exposure. (**E-H**) *wnt2* expression in wild-type and *krm1^nl10^* mutant NMs without exposure to NEO and 4 hours post NEO exposure. Scale bar=20μm. (**I**) Timeline of DMSO or inhibitor exposure, NEO-induced hair cell ablation, and regeneration between 5dpf and 8dpf. (**J-O**) Confocal projects of 8dpf NMs following regeneration, hair cells are labeled with FM1-43 (red) and nuclei are labeled with DAPI (blue) is heterozygous sibling or *krm1^nl10^* NMs with exposure to DMSO, IWR-1, or AZK. (**P**) Quantification of FM1-43-positive cells. DMSO-exposed heterozygous sibling n=26 NM (8 fish) and *krm1^nl10^* n=27 NM (9 fish), IWR-1-exposed heterozygous sibling n=28 NM (7 fish) and *krm1^nl10^* n=17 NM (10 fish), and AZK-exposed heterozygous sibling n=39 NM (7 fish) and *krm1^nl10^* n=24 NM (10 fish). All data presented as mean ±SD, \**p*=024 one-way ANOVA, Tukey’s multiple comparisons test. Scale bar=20μm.

We next sought to determine if altering Wnt activity pharmacologically results in changes in regenerated hair cell numbers in heterozygous sibling and *krm1^nl10^* mutant larvae. Previous work, confirmed by our experiments, demonstrated that regeneration patterns in the lateral line can be altered with exposure to Wnt signaling activators which results in increased hair cell numbers or Wnt inhibitors which leads to fewer hair cells (Fig.4 A-D; ^13^). We sought to determine if we could change the numbers of regenerated hair cells in *krm1^nl10^* mutant larvae though exposure to the Wnt inhibitor IWR-1 or the Wnt activator 1-azakenpaullone (AZK^38^) following NEO-induced hair cell ablation. In larvae exposed to DMSO, we found regeneration of supernumerary FM1-43FX labeled hair cells in *krm1^nl10^* mutant NMs as compared to heterozygous sibling larvae (Fig. 3 J,K,P). This difference is lost in larvae exposed to IWR-1 during regeneration, were we find fewer hair cells forming in *krm1^nl10^* mutant NMs (Fig 3. L,M,P). Reciprocal results were seen when larvae were exposed to AZK during regeneration, resulting in heterozygous sibling NMs forming excess hair cells to levels which were similar to *krm1^nl10^* mutants (Fig. 3, N,O,P). Together these results suggest that loss of Krm1 function results in increased Wnt signaling during regeneration in the posterior lateral line, leading to the formation of supernumerary hair cells. Blocking Wnt signaling with IWR-1 overcomes the bias toward hair cell formation in *krm1^nl10^* mutants, while further activating Wnt with AZK results in increase hair cell regeneration in control larvae, but not mutants.

**Figure 4.**
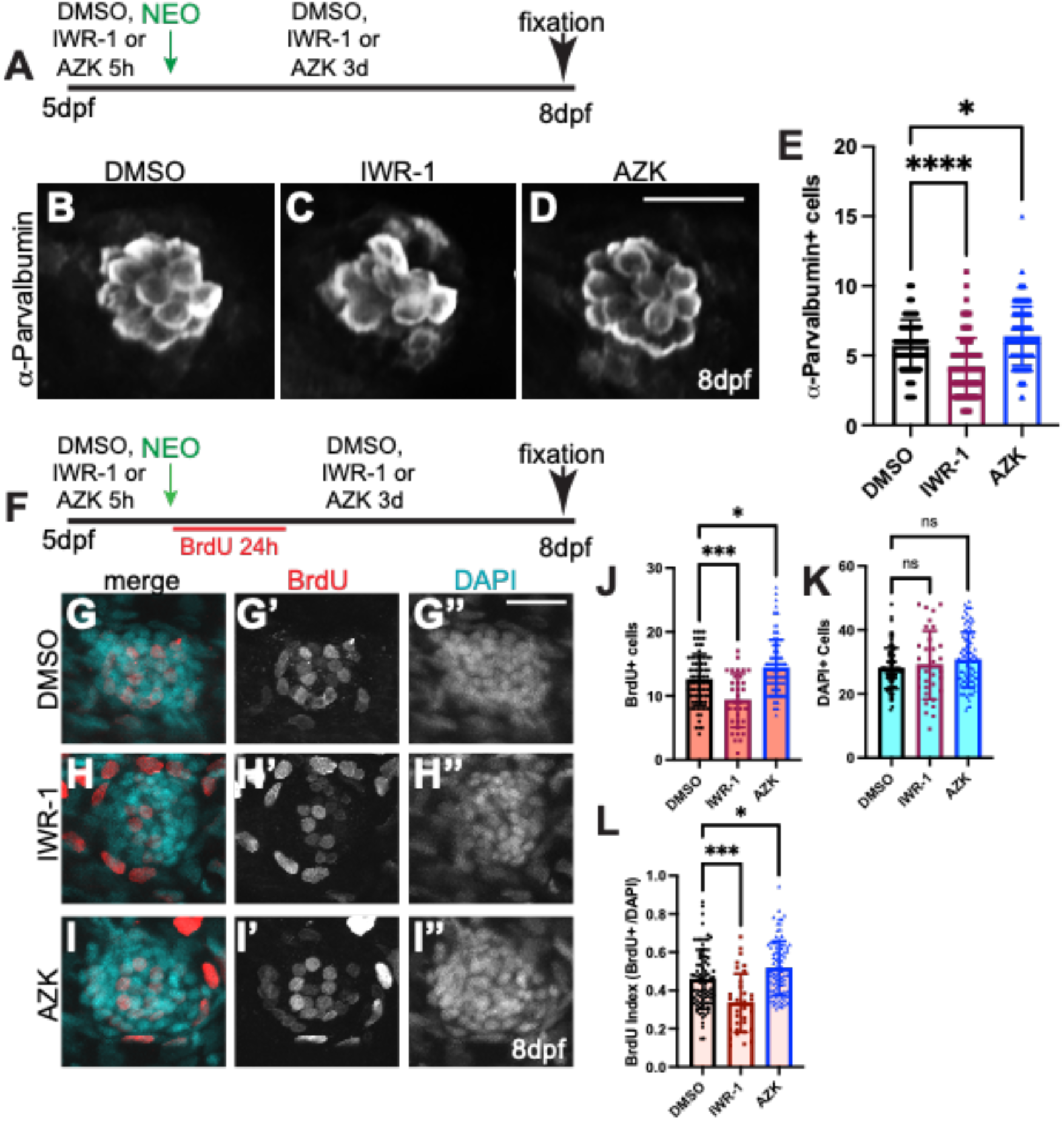
Modulation of the Wnt pathway alters NM regeneration. (**A**) Timeline of DMSO or inhibitor exposure, NEO-induced hair cell ablation, and regeneration between 5dpf and 8dpf. (**C-D**) Confocal projects of 8dpf regenerated hair cells labeled with α-Parvalbumin antibody following exposure to DMSO as a control, IWR-1 to inhibit Wnt signaling, or AZK to activate Wnt activity. (**E**) Quantification of α-Parvalbumin-labeled regenerated hair cells. DMSO n=84 NM (10 fish), IWR-1 n=94 NM (14 fish), and AZK n=156 NM (20 fish). (**F**) Timeline of DMSO or inhibitor exposure, NEO-induced hair cell ablation, BrdU-incorporation for 24h, and regeneration between 5dpf and 8dpf. (**H-I’’**) Confocal projects of 8dpf regenerated hair cells labeled with BrdU to mark proliferating cells and DAPI to label nuclei following exposure to DMSO as a control (**G-G’’**), IWR-1 to inhibit Wnt signaling (**H-H’’**), or AZK to activate Wnt activity (**I-I’’**). (**J-L**) Quantification of BrdU-labeled NM cells following regeneration, DAPI-labeled nuclei, and an index of BrdU-labeled/DAPI-labeled cells. DMSO n=79 NM (14 fish), IWR-1 n=33 NM (5 fish), and AZK n=90 NM (14 fish). All data presented as mean ±SD, \**p*<0.035, \*\*\**p*<0.0007, and **** *p*<0.0001 one-way ANOVA, Tukey’s multiple comparisons test. Scale bar=20μm.

### Regenerative proliferation is not increased in *krm1^nl10^* mutant larvae

Previous work using chemical Wnt inhibitor and overexpression models, suggested that Wnt signaling regulates the proliferation of NM support cells in response to hair cell ablation (^13, 39, 40^). In these studies, inhibition of Wnt signaling using IWR-1 or conditional overexpression of Dkk family members, resulted in a decrease in proliferation during regeneration. Conversely, treatment with Wnt activators LiCl, BIO or Azk, resulted in an increase in proliferation during regeneration (^13, 16, 24–27, 39, 41, 42^). We sought to replicate these experiments by examining BrdU incorporation in DMSO, IWR-1 or AZK exposed larvae during regeneration and found that altering Wnt signaling with these drugs results in a significant decrease in proliferation with IWR-1 (Fig. 4G-G’’’,H-H’’’, J,K,L) and a significant increase in proliferation with AZK during NM regeneration (Fig. 4G-G’’’,I-I’’’, J,K,L).

To determine if loss of Krm1 function alters proliferation in a similar manner during hair cell regeneration, we performed a set of pulse-chase BrdU incorporations during the 3-day period of regeneration following exposure to NEO (Fig. 5A). Cells in regenerating NMs primarily incorporate BrdU during the first day following hair cell ablation (^22^). Following BrdU incorporation during 5-6dpf, we did not find a significant difference in the number of BrdU-positive cells at 8dpf in control or *krm1^nl10^* larvae (Fig. 5B-C’’’, I,J), although when we examined the index of BrdU+ cells compared to total DAPI+ (I) and the index of BrdU+ hair cells (J), we found that there is a significant decrease in *krm1^nl10^* mutants as compared to heterozygous sibling controls. To determine if proliferation is delayed in *krm1^nl10^* during regeneration, we exposed regenerating larvae to BrdU during 6-7dpf (Fig. 5D-E’’’) or 7-8dpf (Fig. 5F-G’’’) following NEO-exposure at 5dpf and then collected the larvae at 8dpf when regeneration was complete. We found that there was no significant change in the total cells incorporating BrdU (Fig. 5 H) or in the index of total cells (Fig. 5I) or hair cells which incorporated BrdU (Fig. 5J). Together these experiments suggest the increase in hair cells present in *krm1^nl10^* mutants following regeneration are not the result of increased proliferation in NM cells.

**Figure 5.**
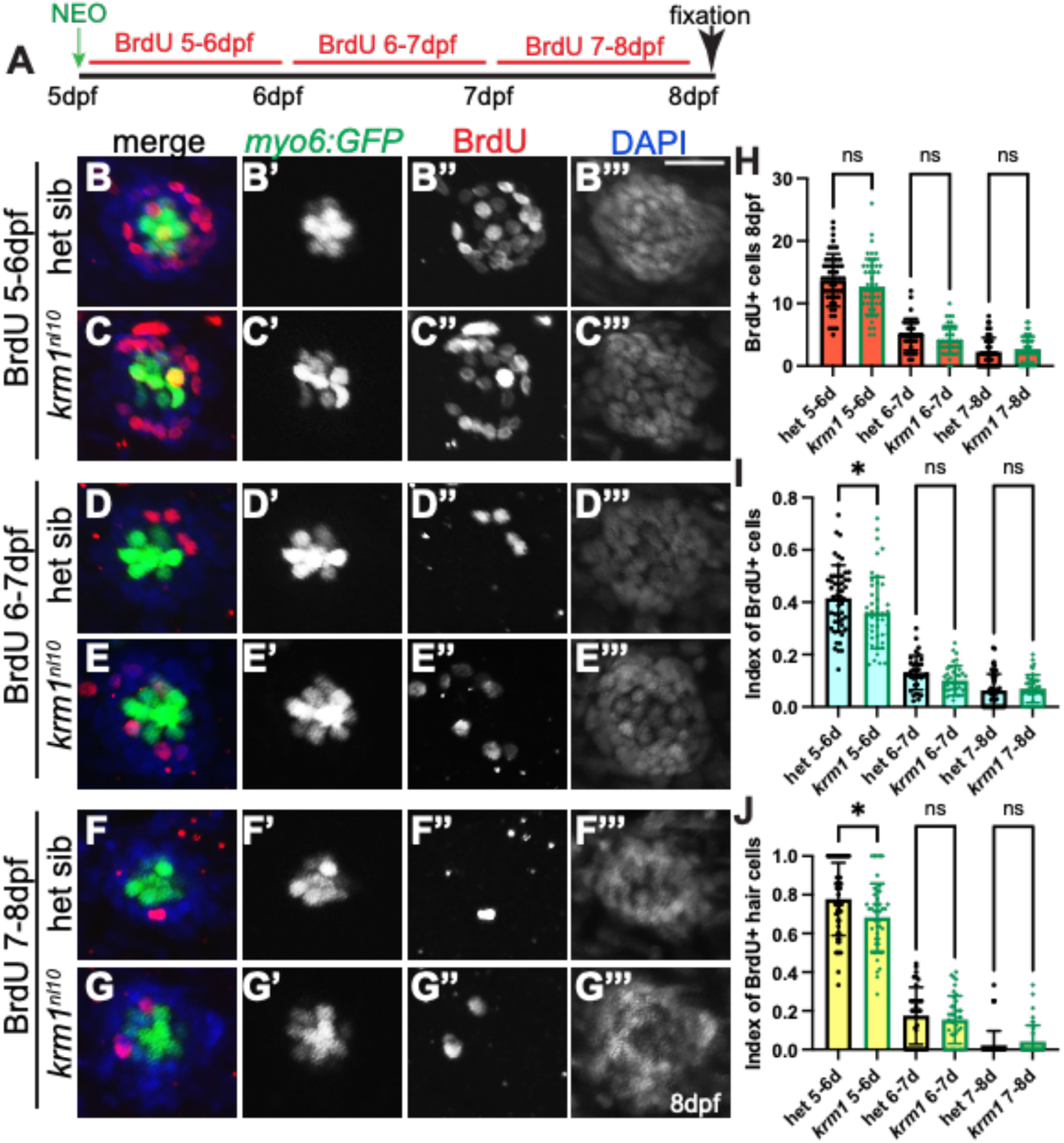
Proliferation does not contribute to the regeneration of supernumerary hair cells in *krm1^nl^*^10^ neuromasts. (**A**) Timeline of neomycin expose at 5dpf, followed by 24 hours of BrdU incubation at 5-6dpf, 6-7dpf, or 7-8dpf, all conditions were allowed to develop until 8dpf and the fixed and processed for imaging. (**B-G’’’** Confocal projections of L2 neuromasts at 8dpf expressing *Tg(myosin6b:GFP)^w186^* to label regenerated hair cells (myo6; green), BrdU incorporation (red), and DAPI labeling of nuclei (blue) at 5-6dpf in heterozygous sibling or *krm1^nl10^* mutant neuromasts NMs, from 6-7dpf,or from 7-8dpf. Scale bar=20μm. (**H-J**) Total numbers of BrdU-positive cells in heterozygous or *krm1^nl10^* mutant neuromasts, index of total BrdU-positive cells (BrdU-labeled cells/DAPI-labeled cells, or Index of BrdU-positive hair cells (BrdU/myo6-labeled cells/myo6-labeled cells). 5-6dpf: het sibling n=50 NMs (12 fish), *krm1^nl10^* n=46 NM (11 fish); 6-7dpf: het sibling n=34 NMs (9 fish), *krm1^nl10^* n=39 NM (9 fish); 7-8dpf: het sibling n=50 NMs (12 fish), *krm1^nl10^* n=40 NM (10 fish). All data presented as mean ±SD, \**p*<0.045 one-way ANOVA, Tukey’s multiple comparisons test.

### Repeated regeneration eliminates supernumerary hair cells in *krm1^nl10^* mutant NMs

The absence of increased proliferation in *krm1^nl10^* mutants leads us to speculate that there may be a population of support cells which are poised to replenish lost hair cells in posterior lateral line NMs which do not rely on entry into the cell cycle. We reasoned that if there is such a population of cells, that repeated regeneration might be sufficient to reduce their number in *krm1^nl10^* mutant larvae and thus eliminating the potential to form supernumerary hair cells (Fig. 6A). We assessed hair cell number after 1 round of regeneration at 8dpf (1x NEO; Fig. 6B-C’, J), 2 rounds of regeneration at 11dp (2x NEO; Fig. 6D-E’, J), 3 rounds of regeneration at 14dpf (3x NEO; Fig. 5F-G’, J), and 4 rounds of 17dpf (4x NEO; Fig. 5H-I’, J). We found following the first round of regeneration, *krm1^nl10^* NMs formed significantly more hair cells compared to heterozygous siblings, as expected from our previous experiments (Fig. 6J), and in each subsequent round of regeneration, we did not find significant differences in regenerated hair cell numbers between mutant and control larvae (Fig. 6J). We also examined to NM cell numbers using DAPI labeling of nuclei and did not find significant differences between heterozygous sibling and *krm1^nl10^* mutant larvae during the repeated rounds of regeneration (Fig. 6B’’-I’’, K), suggesting that there is not a gross change in NM cell number following multiple regeneration events.

**Figure 6.**
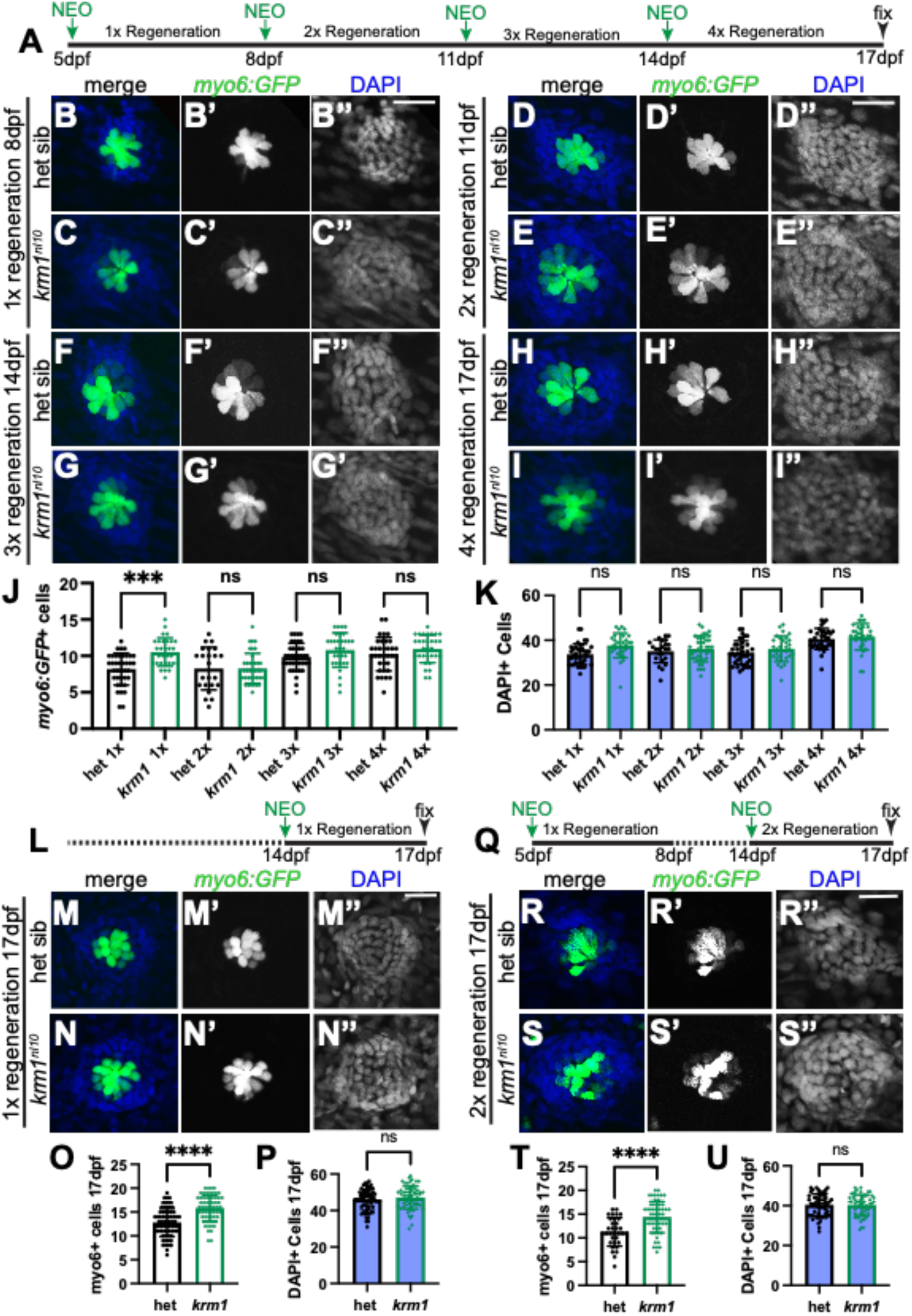
Repeated regeneration eliminates supernumerary hair cells in *krm1^nl^*^10^ larvae. **(A)** Timeline of NEO exposure and repeated regeneration. **(B-I’’)** Confocal projections of NMs expressing *Tg(myosin6b:GFP)^w186^* in hair cells (green) and DAPI-labeling in nuclei following repeated exposures to NEO in heterozygous sibling and *krm1^nl10^* mutant NMs at 8dpf following 1 round of NEO exposure and regeneration, at 11dpf following 2 rounds of NEO exposure and regeneration, NMs at 14dpf following 3 rounds of NEO exposure and regeneration, and at 17dpf following 4 rounds of NEO exposure and regeneration. (**J-K**) Quantification of regenerated *Tg(myosin6b:GFP)^w186^* hair cells or DAPI-labeled nuclei in heterozygous sibling and *krm1^nl10^* mutant NMs at 8dpf, 11dpf, 14dpf, and 17dpf. 8dpf heterozygous sibling n=38 NM (8 fish) and *krm1^nl10^* mutant n=30 NM (10 fish), 11dpf heterozygous sibling n=23 (9 fish) and *krm1^nl10^* n=39 NM (10 fish), 14 dpf heterozygous sibling n=37 NM (9 fish) and *krm1^nl10^* n=31 NM (9 fish), and 17dpf heterozygous sibling n=34 NM (10 fish) and *krm1^nl10^* n=31 NM (9 fish). (**L)** Timeline of a single round NEO-exposure and regeneration between 14-17dpf. **(M-N’’**) Heterozygous sibling and *krm1^nl10^* mutant NMs at 17dpf following 1 round of NEO exposure and regeneration. (**O-P**) Quantification of regenerated *Tg(myosin6b:GFP)^w186^* hair cells and DAPI-positive cells at 17dpf. Heterozygous siblings n=68 NM (12 fish) and *krm1^nl10^* n=62 NM (12 fish). (**Q**) Timeline of 1x NEO exposure at 5dpf and regeneration until 14dpf, 2x NEO from at 14dpf and regeneration until 17dpf. **(R-S’’)** Heterozygous sibling and *krm1^nl10^* mutant NMs at 17dpf following 2x rounds of NEO exposure and regeneration. (**T-U)** Quantification of regenerated *Tg(myosin6b:GFP)^w186^* hair cells and DAPI-positive cells at 17dpf. Heterozygous siblings n=33 NM (8 fish) and *krm1^nl10^* n=58 NM (9 fish). All data presented as mean ±SD, \*\*\**p*=0.0004, **** *p*<0.0001 one-way ANOVA, Tukey’s multiple comparisons test. Scale bar=20μm.

To confirm that the change in regeneration patterns we see with repeated exposure to NEO is based specifically on the response to damage rather than age of the larvae, we exposed 14dpf larvae to NEO and assessed regeneration at 17dpf (Fig. 6L). This single round to regeneration in older larvae resulted in significantly more regenerated hair cells in *krm1^nl10^* mutant NMs (Fig. 6N-N’’’, O,P) as compared to heterozygous siblings (Fig. 6M-M’’’,O,P), indicated that age is not the cause of the absence of supernumerary hair cells following repeated damage. To determine if the poised population of support cells found *krm1^nl10^* can be overwhelmed by regeneration and a prolonged period of recovery, we performed hair cell ablation at 5dpf in heterozygous sibling and *krm1^nl10^* larvae, then allowed them to recover until 14pdf, at which time we performed a second NEO exposure and assessed hair cell regeneration at 17dpf (Fig. 6Q). We found that in contrast to immediately successive rounds of regeneration, a period of recovery re-established the formation of supernumerary hair cells in *krm1^nl10^* mutant NMs (Fig. 6R-U). Together these experiments suggest that there is a population of support cells in *krm1^n^*^10^ mutants that are biased to give rise to hair cells and that these cells can be depleted by repeated regenerations.

### Dorsoventral support cells are increased in *krm1^n^*^10^ mutant neuromasts

Recent studies have found that within lateral line neuromasts, there are distinct populations of support cells that preferentially contribute to hair cell regeneration (^16, 19, 20^). We sought to determine if specific support cell populations are altered in *krm1^nl10^* mutant larvae during development. We began by examining *sost* expression in dorsoventral support cells (Fig. 7A,B) in control and mutant NMs and founda small increase in the *krm1^nl10^* mutants. We quantified *Tg(sost:nlsEos)^w215^*–expressing dorsoventral support cells and *myo6:GFP-*expressing hair cells in heterozygous sibling (Fig. 7C-C’’) and *krm1^nl10^* mutant NMs (Fig. 7C-C’’) at 5dpf and found a significant increase in *sost:nlsEos*+ cells in mutant larvae, including a significant increase in *sost:nlsEos+/myo6:GFP+* hair cells (Fig. 7E). We also examined the *tnfsf10l3-* expressing apical-basal support cells using WISH (Fig. 8A,B) and HCR-FISH and found that there was not significant difference in the number of labeled cells in heterozygous siblings (Fig. 8C-D’’, E) and *krm1^nl10^* mutant NMs (Fig. 8D-D’’’, E). We examined peripheral support cells using *spfr1a* WISH and *Tg(sfpr1a:nlsEos)^w217^* and did not find significant differences between heterozygous and *krm1^nl10^* NMs (Fig. 8F-J) (^19^). Central supports cells were examined using WISH for *six1a*, *lfng,* and *isl1a*, which did not show appreciably different expression patterns in heterozygous larvae and *krm1^nl10^* mutants (Fig. 8K-P) (^15, 16, 20^). These results suggest that loss of Krm1 function during development results in an increase in dorsoventral support cells, as well as the differentiation of hair cells in the posterior lateral line without an increase in overall NM size.

**Figure 7.**
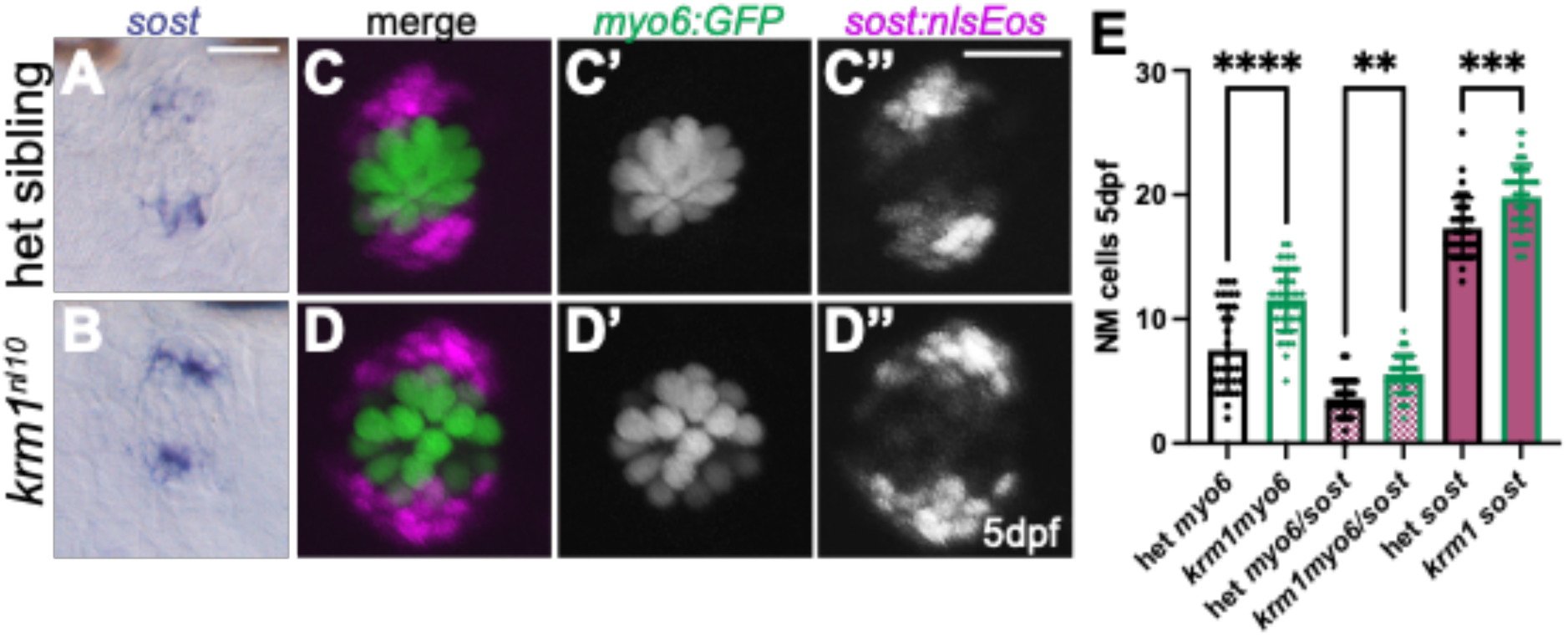
Dorsoventral support cells are increased in *krm1^nl^*^10^ mutant NMs. (**A-B**) RNA in situ hybridization showing *sost* expression in dorsoventral cells in heterozygous sibling and *krm1^nl10^* NMs at 5dpf. (**C-D’’**) Confocal projections of *Tg(myosin6b:GFP)^w186^* labeled hair cells (green) and *Tg(sost:nlsEos)^w215^* labeled dorsoventral support cells (magenta) immediately following photoconversion in live 5dpf heterozygous sibling and *krm1^nl10^* mutant NMs. (**E**) Quantification of *Tg(myosin6b:GFP)^w186^* hair cells, *Tg(myosin6b:GFP)^w186^* and *Tg(sost:nlsEos)^w215^* expressing hair cells, and total *Tg(sost:nlsEos)^w215^* expressing NM cells. Heterozygous siblings n=34 NM (8 fish) and *krm1^nl10^* n=46 NM (9 fish). All data presented as mean ±SD, \*\**p*=0.006, \*\*\**p*=0.0001, and **** *p*<0.0001 one-way ANOVA, Tukey’s multiple comparisons test. Scale bar=20μm.

**Figure 8.**
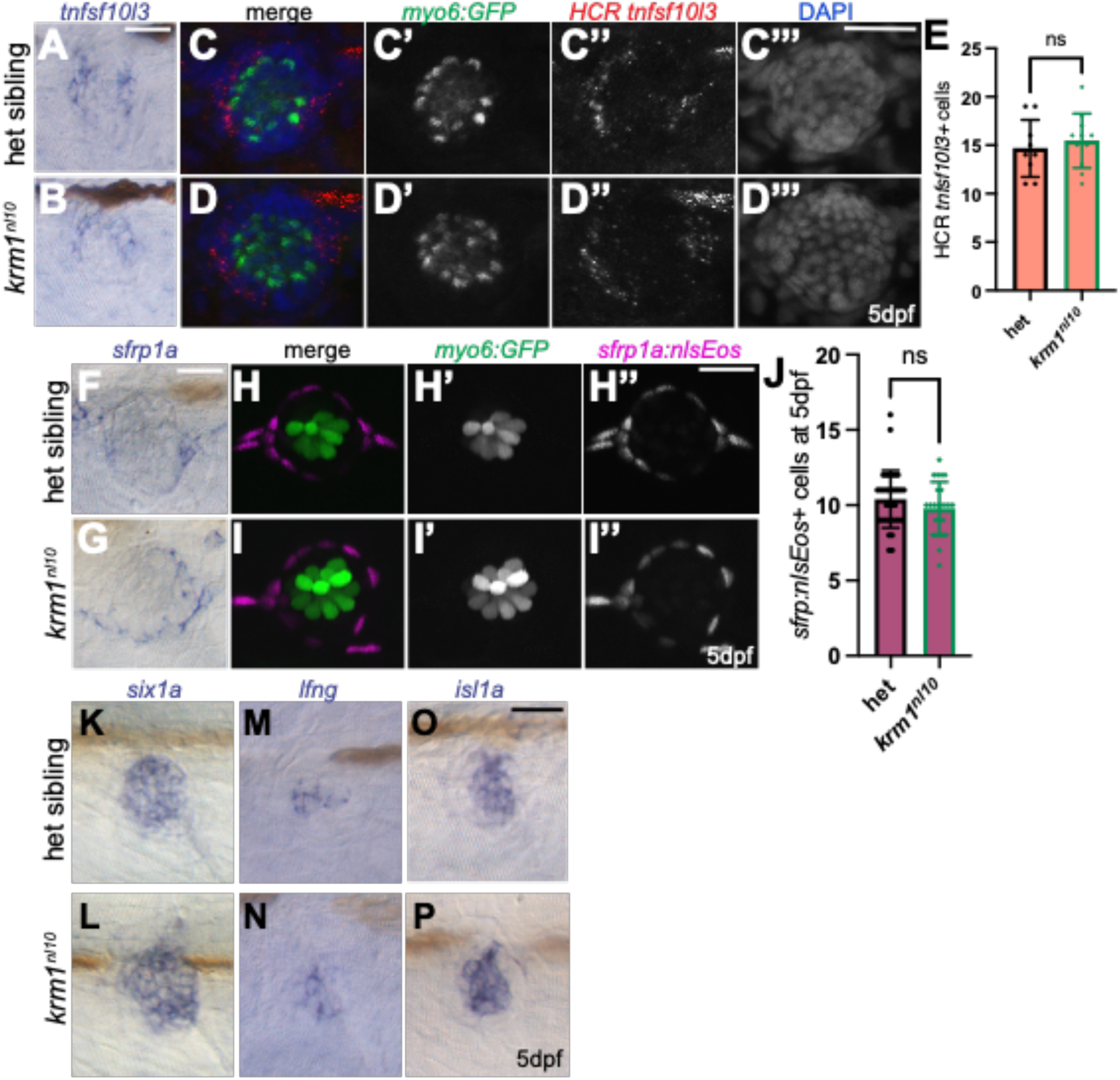
Anterior/posterior and peripheral support cells are not significantly altered in *krm1^nl^*^10^ NMs. **(A-B)** RNA in situ hybridization showing *tnfsd10l3* expression in anterior-posterior cells in heterozygous sibling and *krm1^nl10^* NMs at 5dpf. (**C-D’’’)** Confocal projections of HCR-FISH showing *tnfsf10l3* expression (red) in 5dpf *Tg(myosin6b:GFP)^w186^* larvae (green) with nuclei labeled with DAPI (blue) in heterozygous sibling and *krm1^nl10^* mutant NMs. (**E)** Quantification of HCR *tnfsf10l3*-labeled cells at 5dpf, heterozygous siblings n=9 NM (9 fish) and *krm1^nl10^* n=11 (11 fish). (**F-G)** RNA in situ hybridization showing *sfrp1a* expression in anterior-posterior cells in heterozygous sibling and *krm1^nl10^* NMs at 5dpf. (**H-I’’)** Live confocal projections of *Tg(myosin6b:GFP)^w186^* larvae (green) with nuclei labeled with DAPI (blue). Confocal projections of *Tg(myosin6b:GFP)^w186^* labeled hair cells (green) and *Tg(sfrp1a:nlsEos)^w217^* labeled dorsoventral support cells (magenta) immediately following photoconversion in live 5dpf heterozygous sibling and *krm1^nl10^* mutant NMs. (**J)** Quantification of total *Tg(sost:nlsEos)^w215^* expressing NM cells. Heterozygous siblings n=35 NM (10 fish) and *krm1^nl10^* n=24 NM (10 fish). (**K-P)** RNA in situ hybridization for central support cell markers in heterozygous sibling ( *krm1^nl10^* NMs showing expression of *six1a*, *lunatic fringe* (*lfng*), and *islet1a* (*isl1a*) at 5dpf. All data presented as mean ±SD, one-way ANOVA, Tukey’s multiple comparisons test. Scale bar=20μm.

### Dorsoventral cell contribute to the regeneration of supernumerary hair cells in *krm1^nl10^* NMs

To analyze the contribution of *sost:nlsEos*+ support cells to hair cell regeneration we photoconverted larvae carrying the *Tg(sost:nlsEos)^w215^* and *Tg(myo6b:GFP)^w186^* transgenes by exposing them to UV light at 5dpf and assessing cell numbers at 8dpf in live larvae. Under homeostatic conditions (Fig. 9A), without exposure to NEO, we find significantly more *myo6:GFP*+ hair cells and *sost:nlsEos*+ in *krm1^nl10^* mutant NMs (Fig. 9C-C’’, D) as compared to heterozygous siblings (Fig. 9B-B’’, D). We did not find a significant change in the hair cells co-expressing *myo6:GFP* and *sost:nlsEos* (Fig.9D), suggesting that under homeostatic conditions, few hair cells arise between 5-8dpf. To assess the contribution of *sost:nlsEos*+ dorsoventral support cells, we exposed larvae to NEO following photoconversion at 5dpf and allowed regeneration to progress to 8dpf before live imaging (Fig. 9E). Under regeneration conditions we found that there was a significant increase in *myo:GFP*+ hair cells, hair cells expressing *myo:GFP* and *sost:nlsEos*, and *sost:nlsEos*+ cells in *krm1^nl10^* mutant NMs (G-G’’, H) as compared to heterozygous control larvae (Fig. 9F-F’’, H). These results suggest that the supernumerary *sost:nlsEos+* support cells in *krm1^nl10^* mutant give rise to excess hair cells during regeneration. We next asked if repeated regeneration could overwhelm *sost:nlsEos+* dorsoventral support cells and reduce the number of regenerated hair cells. For these experiments, we performed photoconversion and NEO exposure on *sost:nlsEos* and *myo6:GFP* larvae at 5dpf and again at 8dpf, allowing the fish to undergo two rounds of regeneration before collection and live imaging at 11dpf (Fig. 9I). In these experiments, there was no longer a significant difference in the total number of regenerated *myo:GFP+* hair cells, hair cells co-expressing *myo6:GFP* and *sost:nlsEos*, or total *sost:nlsEos+* support cells between heterozygous siblings (Fig. 9J-J’’,L) and *krml^nl10^* mutant larvae (Fig. 9K-K’’, L). These results suggest that dorsoventral support cells are the source of supernumerary hair cells in *krm1^nl10^* mutants.

**Figure 9.**
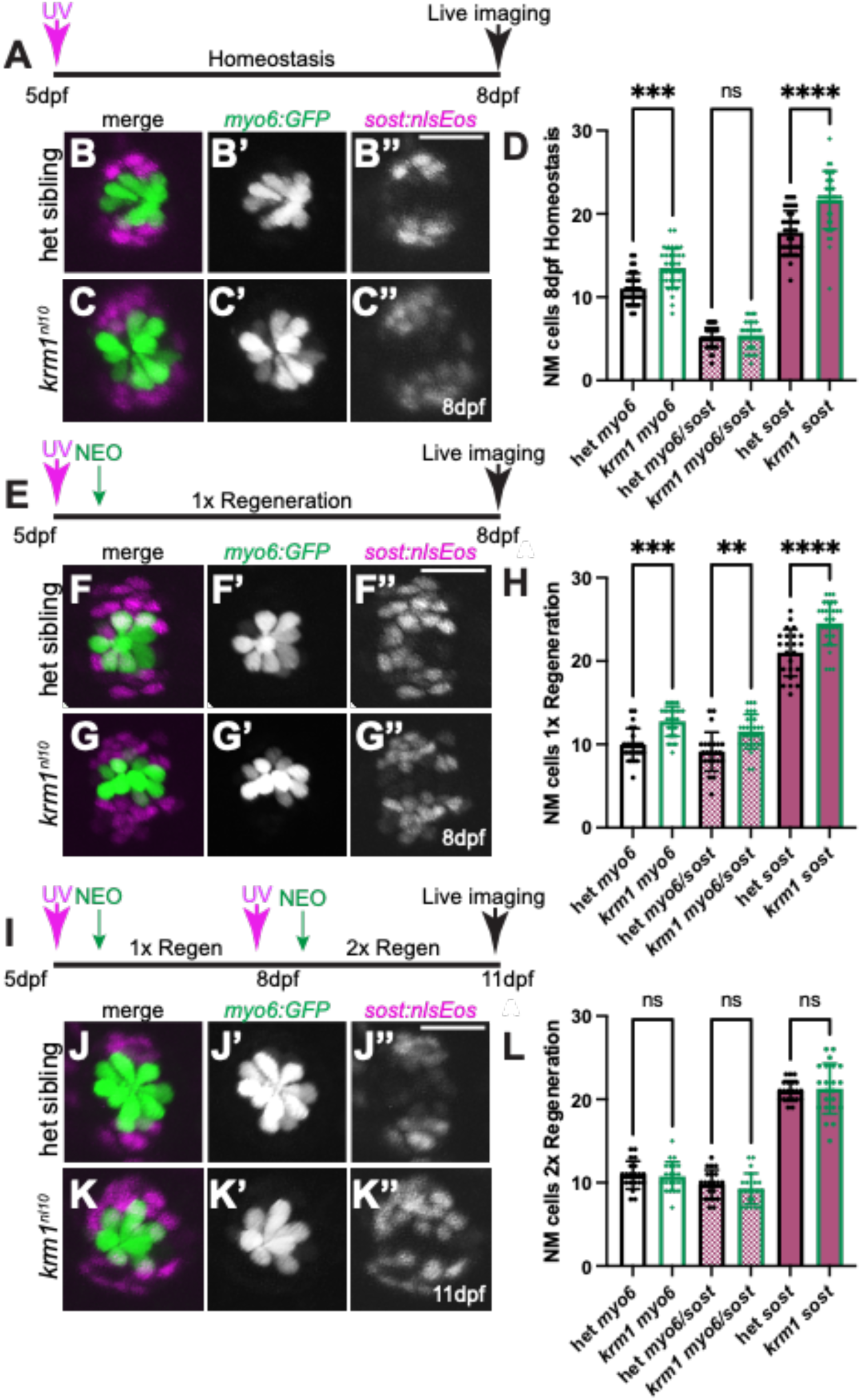
Dorsoventral support cells contribute to supernumerary hair cells following regeneration in *krm1^nl^*^10^ mutant NMs. (**A**) Timeline for *Tg(sost:nlsEos)^w215^* photoconversion at 5dpf and homeostatic development until 8dpf. (**B-C’’**) Homeostasis at 8d of *Tg(myosin6b:GFP)^w186^* labeled hair cells (green) and *Tg(sost:nlsEos)^w215^* labeled dorsoventral support cells (magenta) following photoconversion in 5dpf heterozygous sibling and *krm1^nl10^* mutant NMs. (**D**) Quantification of *Tg(myosin6b:GFP)^w186^* hair cells, *Tg(myosin6b:GFP)^w186^* and *Tg(sost:nlsEos)^w215^* expressing hair cells, and total *Tg(sost:nlsEos)^w215^* expressing NM cells. Heterozygous siblings n=21 NM (12 fish) and *krm1^nl10^* n=22 NM (12 fish). (**E**) Timeline for *Tg(sost:nlsEos)^w215^* photoconversion and NEO at 5dpf and 1 round of regeneration until 8dpf. (**F-G’’**) 1x regeneration of *Tg(myosin6b:GFP)^w186^* labeled hair cells (green) and *Tg(sost:nlsEos)^w215^* labeled dorsoventral support cells (magenta) following photoconversion at 5dpf and NEO-exposure in heterozygous sibling and *krm1^nl10^* mutant NMs. (**H**) Quantification of *Tg(myosin6b:GFP)^w186^* hair cells, *Tg(myosin6b:GFP)^w186^* and *Tg(sost:nlsEos)^w215^* expressing hair cells following 1x regeneration, and total *Tg(sost:nlsEos)^w215^* expressing NM cells. Heterozygous siblings n=23 NM (8 fish) and *krm1^nl10^* n=31 NM (8 fish). (**I**) Timeline for *Tg(sost:nlsEos)^w215^* photoconversion and NEO at 5dpf and again at 8dpf, for 2 rounds of regeneration until 11dpf. (**J-K’’**) 2x regeneration of *Tg(myosin6b:GFP)^w186^* labeled hair cells (green) and *Tg(sost:nlsEos)^w215^* labeled dorsoventral support cells (magenta) following photoconversion and NEO-exposure at 5dpf and 8dfp in heterozygous sibling and *krm1^nl10^* mutant NMs. (**L**) Quantification of *Tg(myosin6b:GFP)^w186^* hair cells, *Tg(myosin6b:GFP)^w186^* and *Tg(sost:nlsEos)^w215^* expressing hair cells following 2x regeneration, and total *Tg(sost:nlsEos)^w215^* expressing NM cells. Heterozygous siblings n=21 NM (8 fish) and *krm1^nl10^* n=22 NM (7 fish). All data presented as mean ±SD, \*\**p*=0.002, \*\*\**p*=0.0002, and **** *p*<0.0001 one-way ANOVA, Tukey’s multiple comparisons test. Scale bar=20μm.

### Notch inhibition alters cell numbers in *krm1^nl10^* NMs

The Notch pathway is critical for regulating hair cell regeneration and inhibition of Notch signaling greatly increases the number of hair cells that regrow following damage (^16, 19, 22, 24^). Specifically, Notch signaling regulates proliferation of NM cells; inhibition of the Notch pathway results in increased support cell proliferation, while activating Notch signaling decreased proliferation (^16, 24^). As we do not see a significant change in proliferation during regeneration in our *krm1^nl10^* mutant line, we reasoned that the Notch pathway might play more central regulatory in this process as compared to Wnt signaling. To test this, we exposed heterozygous sibling and *krm1^nl10^* mutant 5pdf larvae to DMSO or the ψ-secretase inhibitor LY441575 for 5 hours, then ablated hair cells with NEO, we then returned the larvae to DMSO or LY441575 that included BrdU for 24h, and then we incubate larvae for 2 more days in DMSO or LY441575 (Fig. 10A). We found that hair cells and BrdU incorporation significantly increased in both heterozygous sibling (Fig. 10D-D’’’, F,G) and *krm1^nl10^* NMs (Fig. 10E-E’’’, FG) when treated with LY441575, as compared to DMSO treated larvae (Fig. 10A-B’’’, F,G). Though the number of cells incorporating BrdU increased when Notch was inhibited, there was no significant difference between heterozygous siblings and *krm1^nl10^* mutant larvae (Fig. 10G), suggesting that proliferation is regulated independently from Krm1 function.

**Figure 10.**
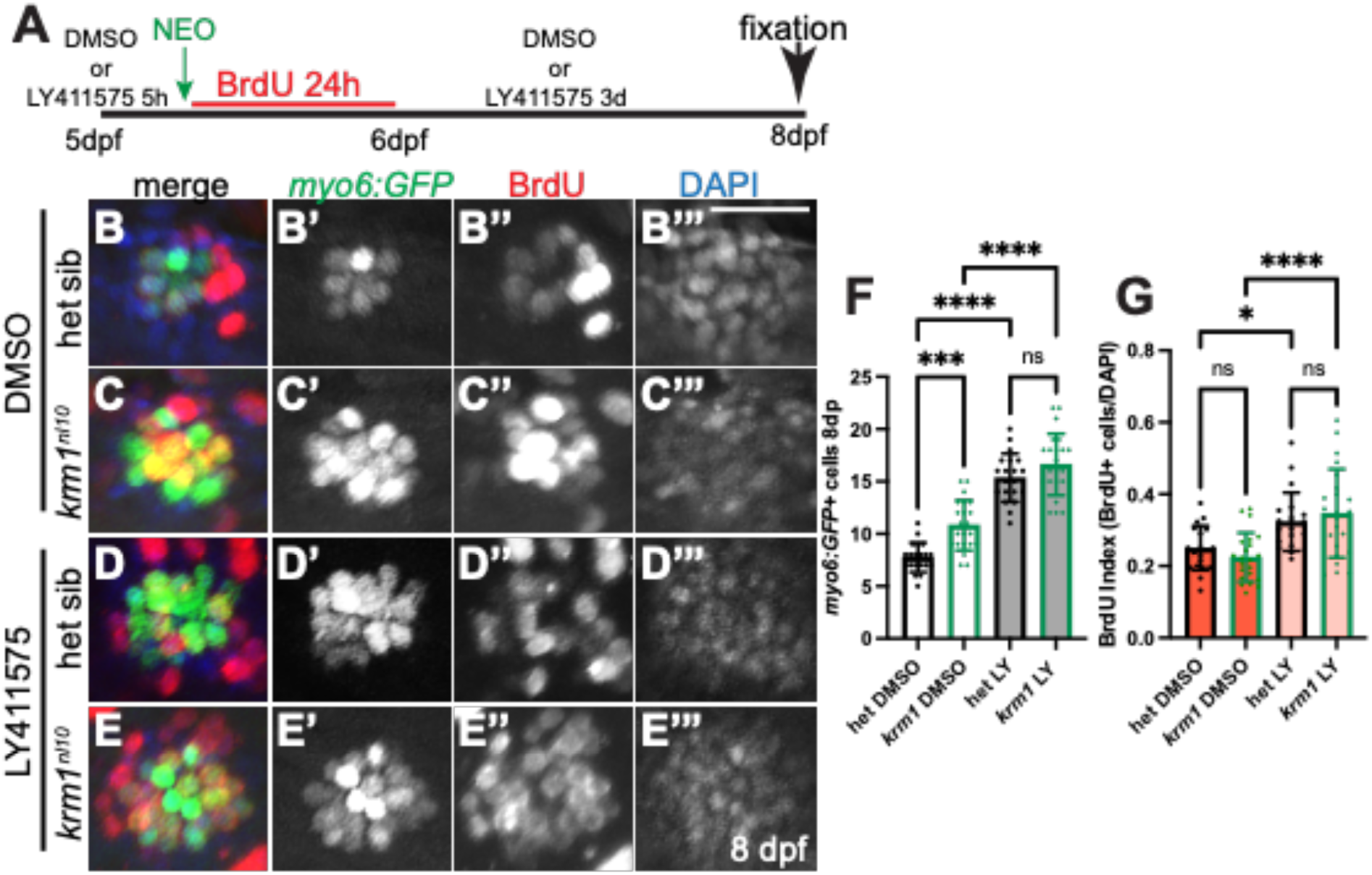
Notch signaling regulates cellular proliferation during regeneration in *krm1^nl^*^10^ neuromasts. (**A**) Timeline of in DMSO or LY411575 for 5-hours at 5dpf, NEO exposure, BrdU incubation for 24h hours in the presence of DMSO or LY411575, and then regeneration to 8dpf with exposure to DMSO or LY411575. (**B-B’’’**) Confocal projections of L2 neuromasts at 8dpf expressing *Tg(myosin6b:GFP)^w186^* to label regenerated hair cells (*myo6:GFP*; green), BrdU incorporation (red), and DAPI labeling of nuclei (blue) in heterozygous sibling and *krm1^nl10^* mutant larvae following exposure to DMSO or LY411575. (**F-G**) Quantification of *Tg(myosin6b:GFP)^w186^* labeled hair cells (green) cells at 8dfp following regeneration during exposure to DMSO or LY411575 and index of BrdU-positive cells/DAPI-labeled nuclei in NMs. DMSO-exposed heterozygous siblings n=20 NM (8 fish) and *krm1^nl10^* n=22 NM (8 fish); LY411575-exposed heterozygous siblings n=20 NM (8 fish) and *krm1^nl10^* n=23 NM (8 fish). All data presented as mean ±SD, \**p*<0.05, \*\*\**p*=0.0004, and **** *p*<0.0001 one-way ANOVA, Tukey’s multiple comparisons test. Scale bar=20μm.

Previous work demonstrated that both the dorsoventral and anterior-posterior populations of support cells increase in number when the Notch pathways in inhibited during regeneration (^19^). Since the Notch pathway regulates the regeneration of both hair cells and dorsoventral support cells, we decided to assess what happens to these cell populations in *krm1^nl10^* mutant larvae when Notch signaling was conditionally inhibited during regeneration. RNA in situ hybridization shows similar expression of Notch pathways members *notch1a*, *notch3*, *deltaA*, and *deltaD*, as well as the hair cell progenitor marker *atoh1a,* in heterozygous sibling and *krm1^nl10^* mutant prior to NEO-exposure at 5dpf and 1d-post hair cell ablation (Fig. 11A-T). To assess the role of dorsoventral support cells during regeneration exposed heterozygous or *krm1^nl10^* mutant larvae carrying the *Tg(sost:nlsEos)^w215^* and *Tg(myo6b:GFP)^w186^* transgenes to DMSO or LY441575. At 5dpf we incubated the larvae in DMSO or inhibitor for 5 hours prior to photoconversion with a UV light and the exposure to NEO to ablate hair cells, the larvae were then allowed to regenerate for 3 days and imaged live at 8dpf (Fig. 12A). In agreement with our previous experiments, we found that *krm1^nl10^* mutant larvae exposed to DMSO showed a significant increase in the number of regenerated hair cells and dorsoventral *sost:nlsEos*+ support cells (Fig. 12C-C’’, F) as compared to heterozygous sibling controls (Fig. 12B-B’’, F). In comparison to DMSO conditions, LY441575 treatment resulted in a significant increase in the regenerated hair cells in both heterozygous (Fig. 12D-D’, F) and *krm1^nl10^* mutant NMs (Fig. 12E-E’,F), eliminating the difference in populations we see in previous experiments (Fig. 12F). When we examined *sost:nlsEos+* support cells in larvae exposed to LY441575, we found a significant increase in heterozygous NMs as compared to DMSO conditions (Fig. 12B’’, D’’, F), but not in *krm1^nl10^* mutant larvae (Fig. 12C’’, E’’, F). However, even following LY441575 exposure, *krm1^nl10^* mutant larvae had significantly more *sost:nlsEos+* cells as compared to LY441575-treated heterozygous controls (Fig. 12F).

**Figure 11.**
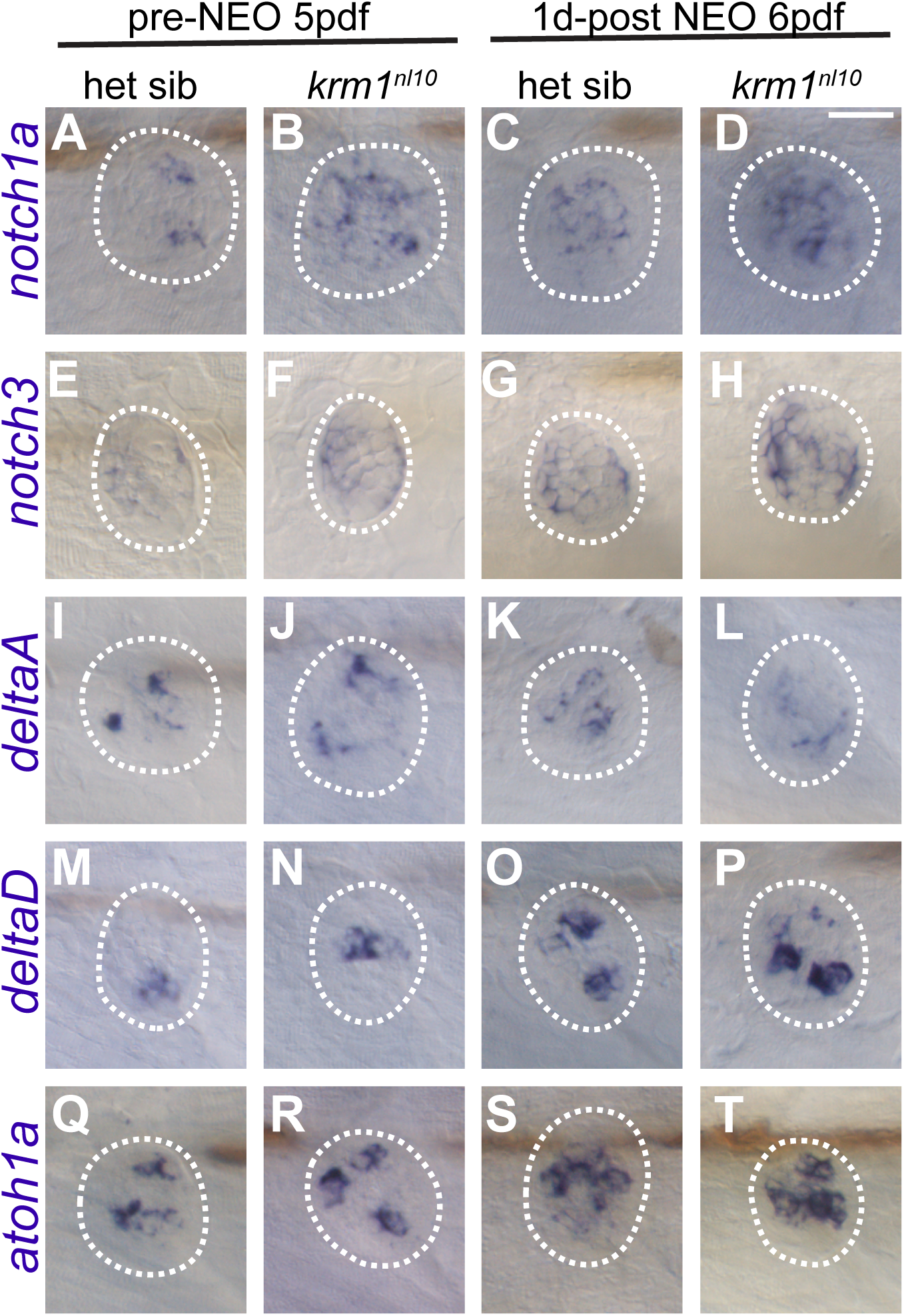
RNA in situ hybridization for Notch pathway members during regeneration. RNA in situ hybridization for Notch pathway members in NMs. Expression of *notch1a* and *notch3* in heterozygous sibling (**A,E**) and *krm1^nl10^* (**B,F**) NMs at 5dpf and increased 1-day poster NEO exposure in heterozygous sibling (**I,M**) and *krm1^nl10^* (**J,N**) NMs at 6dpf. Expression of *deltaA* and *deltaD* in heterozygous sibling (**A,E**) and *krm1^nl10^* (**B,F**) NMs at 5dpf and increased 1-day poster NEO exposure in heterozygous sibling (**K,O**) and *krm1^nl10^* (**L,P**) NMs at 6dpf. The hair cell precursor marker *atoh1a* is expressed at moderate levels in heterozygous sibling (**Q**) and *krm1^nl10^* (**R**) NMs at 5dpf and increased 1-day poster NEO exposure in heterozygous sibling (**S**) and *krm1^nl10^* (**T**) NMs at 6dpf. Scale bar=20μm.

**Figure 12.**
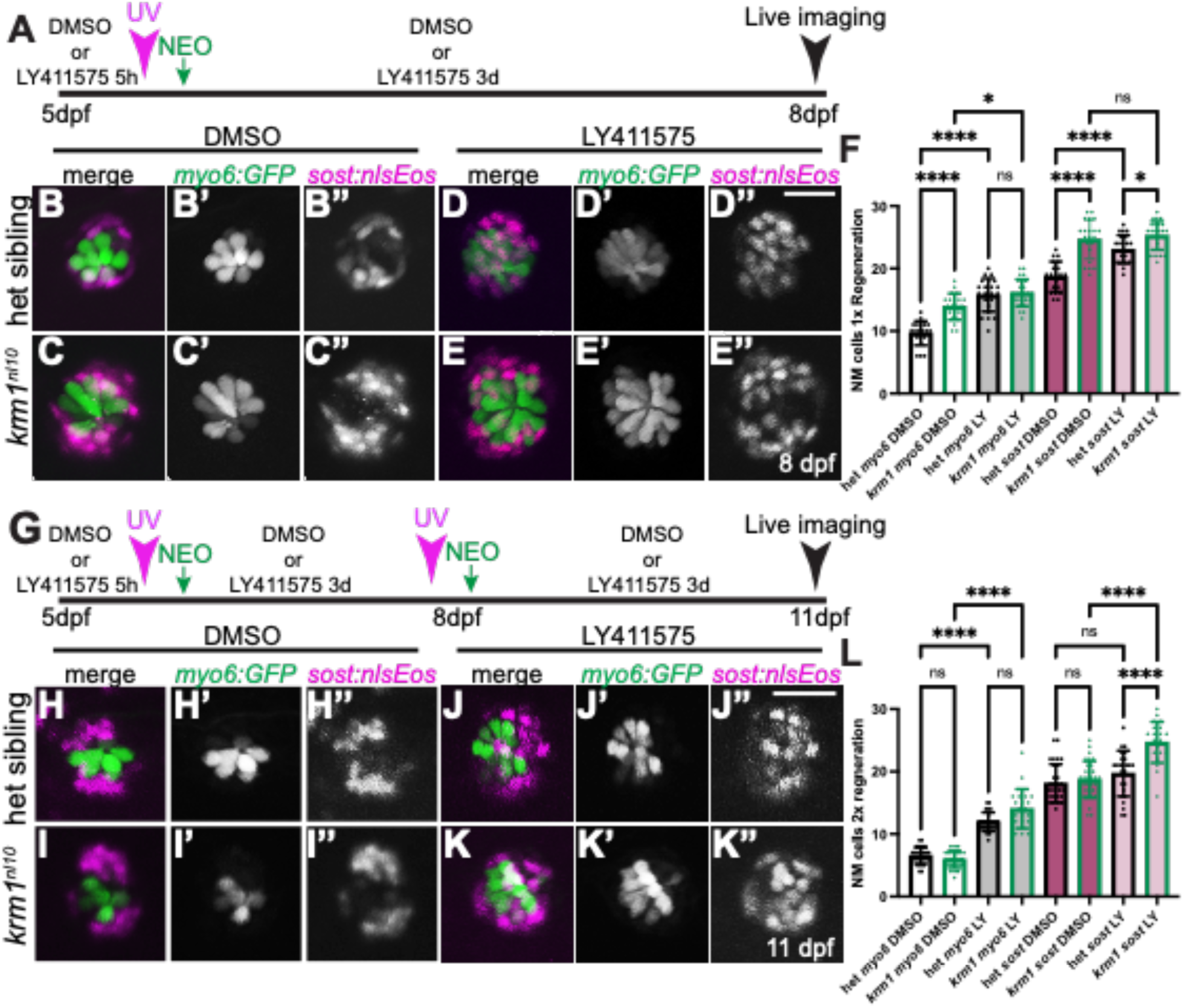
Notch signaling alters NM cell numbers following repeated damage. (**A**) Timeline of drug exposure from 5-8dpf, photoconversion, NEO exposure, regeneration, and live imaging. (**B-E’’**) Live confocal projections of *Tg(myosin6b:GFP)^w186^* labeled hair cells (green) and *Tg(sost:nlsEos)^w215^* labeled dorsoventral support cells (magenta) in larvae exposed to DMSO or LY411575 and 1x regeneration in heterozygous sibling *krm1^nl10^* mutant larvae. (**F**) Quantification of *Tg(myosin6b:GFP)^w186^* labeled hair cells (green) and *Tg(sost:nlsEos)^w215^* labeled cells at 8dfp following regeneration during exposure to DMSO or LY411575. DMSO-exposed heterozygous siblings n=20 NM (9 fish) and *krm1^nl10^* n=21 NM (9 fish); LY411575-exposed heterozygous siblings n=22 NM (9 fish) and *krm1^nl10^* n=25 NM (11 fish). (**G**) Timeline for drug exposure, *Tg(sost:nlsEos)^w215^* photoconversion and NEO at 5dpf and again at 8dpf, for 2 rounds of regeneration until 11dpf. (**H-K’’**) Heterozygous sibling and *krm1^nl10^* mutant larvae exposed to DMSO or LY411575 during 2x regeneration. (**L**) Quantification of *Tg(myosin6b:GFP)^w186^* hair cells, *Tg(myosin6b:GFP)^w186^* and *Tg(sost:nlsEos)^w215^* expressing hair cells following 2x rounds of regeneration in the presence of DMSO or LY411575. DMSO-exposed heterozygous siblings n=38 NM (16 fish) and *krm1^nl10^* n=39 NM (14 fish); LY411575-exposed heterozygous siblings n=25 NM (10 fish) and *krm1^nl10^* n=24 NM (9 fish). All data presented as mean ±SD, \**p*<0.04 and **** *p*<0.0001 one-way ANOVA, Tukey’s multiple comparisons test. Scale bar=20μm.

As inhibition of Notch signaling increases proliferation, dorsoventral support cell numbers, and hair cell regeneration in *krm1^nl10^* mutant larvae, we next asked if this increase was sufficient to overcome the depletion of poised support cells we found following multiple rounds of regeneration in mutant NMs. We performed two successive rounds of regeneration between 5dpf and 11dpf using heterozygous sibling and *krm1^nl10^ myo6:GF)/sost:nlsEos*-expressing larvae is continuous exposure to DMSO or LY441575 (Fig. 12G). We found in our DMSO treated conditions that 2 rounds of regeneration was sufficient to eliminate supernumerary *sost:nlsEos*+ support cells and *myo:GFP*+ hair cells in *krm1^nl10^* mutant larvae as compared to heterozygous controls (Fig. 12H-I’’,L). In contrast, inhibition of Notch signaling resulted in a significant increase in both *sost:nlsEos*+ and *myo:GFP*+ cells in heterozygous sibling and *krm1^nl10^* mutant NMs following successive rounds of NEO exposure (Fig. 12J-K’’,L). Together these results suggest that the Notch pathway regulates support cell proliferation and hair cell regeneration thought mechanisms which are distinct from those regulated by the Wnt pathway and Kremen1 function.

## Discussion

Our work demonstrates that Krm1 functions in the zebrafish posterior lateral line to regulate the number of dorsoventral support cells and hair cells during NM development and regeneration. Under most conditions Krm1 acts to inhibit canonical Wnt signaling, and the results reported here agree with this function in the lateral line(^29, 31^). We found that in contrast to previous analysis of Wnt signaling in the lateral line, regeneration of supernumerary hair cells in *krm1^nl10^* mutants occurs through the direct differentiation of support cells without an increase in proliferation (^13, 16, 24–27, 39, 41, 42^).

Our analysis of Krm1 function during regeneration in the zebrafish posterior lateral line revealed that there is a population of support cells that is poised to give rise to new hair cells even in the absence of cellular proliferation. These support cells are members of the *sost:nlsEos*-positive dorsoventral population, which has been shown to be the primary source of regenerating hair cells following neomycin induced damage (^19^). As Krm1 functions to inhibit the canonical Wnt pathway, our results examining the loss of Krm1 function suggest that Wnt signaling acts to regulate the number of poised progenitor cells. We find that with repeated ablation of hair cells and successive rounds of regeneration, we can deplete these poised progenitor cells and eliminate supernumerary hair cells in *krm1^nl10^* mutants.

### Specific function of support cell populations during lateral line regeneration

Recent studies have begun to identify subsets of support cells within the zebrafish lateral line which seem to play specific roles during regeneration and have distinct genetic profiles (^15, 16, 19, 20^). Within these specialized support cells there are populations that primarily contribute to self-renewal and others that will give rise to regenerating hair cells. Although *krm1* is expressed throughout the NM, loss of Krm1 function seems to particularly result in an increase in the dorsoventral population of *sost:nlsEos+* support cells and mature hair cells. Within these cells, there seems be to a further specialized population of cells which can differentiate into hair cells without first undergoing proliferation. As we find an increase in these cells in *krm1^nl10^* mutants, it seems that Wnt signaling regulates their numbers. As we develop more precise expression profiles of cells participating in NM regeneration, we will hopefully be able to specifically determine the identity of these poised support cells.

### Wnt and Notch signaling in the lateral line

Previous studies have demonstrated that members of the Wnt pathway are expressed at low levels in the dorsal ventral poles of the NMs during homeostasis and are strongly upregulated beginning at 3-hours post hair cell ablation with neomycin (^15, 16, 19, 24^). Several studies have used pharmacological manipulation of the Wnt pathway to dissect its function during regeneration of the zebrafish lateral line (^16, 24, 25, 42, 43^). These studies, as well as our work, suggest that altering Wnt activity specifically results in changes in support cells proliferation; activating Wnt with LiCl, BIO or AZK results in increased proliferation and inhibition of Wnt signaling with IWR-1 leads to decreased support cell proliferation (Fig. 4; ^16, 24, 25, 42, 43^). With this history, we were surprised to find that we did not find an increase in proliferation in our *krm1^nl10^* mutants during regeneration and in fact saw a decrease in the proportion of BrdU-positive support cells during regeneration (Fig. 5).

The difference between pharmacological manipulation and the *krm1^nl10^* mutants might be a result of the specificity of the targets affected. AZK and BIO both activated the Wnt pathway though inhibition of glycogen synthase kinase 3β (GSK3β) and disruption of the destruction complex, releasing β-catenin and allows Wnt-mediated gene transcription (^38^). Though GSK3β is strongly associated with canonical Wnt signaling, it also regulates cell behaviors via multiple pathways, in particular through the AKT/mTOR pathway to promote proliferation and regeneration (^44^). Along similar lines, IWR-1 inhibits Tankyrase, which results in the stabilization of Axin and the increased phosphorylation and degradation of β-catenin. Tankyrase plays roles in many signaling pathways that can impinge on cellular proliferation (^45^). Thus, the changes in proliferation seem during regeneration with exposure to these pharmacological manipulations may be the results of alterations in pathways in addition to canonical Wnt signaling.

The Notch/Delta pathway is critical for regulating cellular behavior in during development and regeneration in the lateral line (^16, 19, 22, 24, 27^). Inhibition of Notch signaling results in dramatic increases in proliferation, support cell numbers, and hair cell formation (^16, 22, 24^). Notch has been shown to act upstream of Wnt signaling during the process of regeneration, though previous work examining the results of simultaneously manipulating the Notch and Wnt pathways results in changes in the levels of proliferation during lateral line regeneration (^16, 24^). In contrast, our experiments assessing regeneration using the *krm1^nl10^* mutant line, suggest that upregulating Wnt activity regulation hair cell progenitor specification and differentiation, while inhibiting Notch signaling increases proliferation and increased hair cell differentiation. The concurrent upregulation of Wnt and inhibition of Notch results in a proportionally smaller increase in dorsoventral support cells and hair cells compared to changes in heterozygous siblings, suggesting the Wnt signaling is predominantly responsible for cell identity in lateral line neuromasts, while Notch signaling regulates support cell proliferation. Future work examining additional Wnt pathway mutants should provide a more detailed understanding of the role of Wnt in regulating regeneration.

### Krm1 and human hearing

Recent studies using mammalian inner ear cell cultures suggest that manipulating the Notch and Wnt pathways are critical to regrowing damaged hair cell in this model (^46, 47^). Interestingly, there is conflicting results for upregulation and down regulation of Notch signaling as a mechanism to promote regeneration (^46, 47^), while upregulation of Wnt signaling is consistently necessary. Krm1 is expressed in the support cells of the mammalian cochlea, making it a potential target for therapeutic intervention, in which inhibition of Krm1 function might be required for support cells to re-enter a more progenitor-like stage and to then differentiate as mechanosensory hair cells to restore hearing. Future work will be needed to determine if regulation of Krm1 function also plays a role in hair regeneration in the inner ear.

### Limitations of the study

In this study, we used the *krm1^nl10^* mutant zebrafish line to determine if Krm1 function regulates hair cell regeneration. We determined that *krm1^nl10^* mutants form more dorsoventral support cells in their neuromasts as compared to controls, and that these cells can regenerate supernumerary hair cells in the absence of increase proliferation. One of the primary limitations of this study is that we do not know the precise identity of these poised dorsoventral cells. Future studies examining existing gene expression profiles of the lateral line or single-cell RNA sequencing of *krm1^nl10^* mutants, will help identify this poised progenitor population. Another limitation is that the current study does not address the role of these poised progenitor cells in wild-type fish under homeostatic conditions. It is possible that this is a small population of support cells, which are able to rapidly replenish hair cells damaged during the normal life cycle of the fish. Future work is needed to address these questions.

## Materials and Methods

**Table 1:**
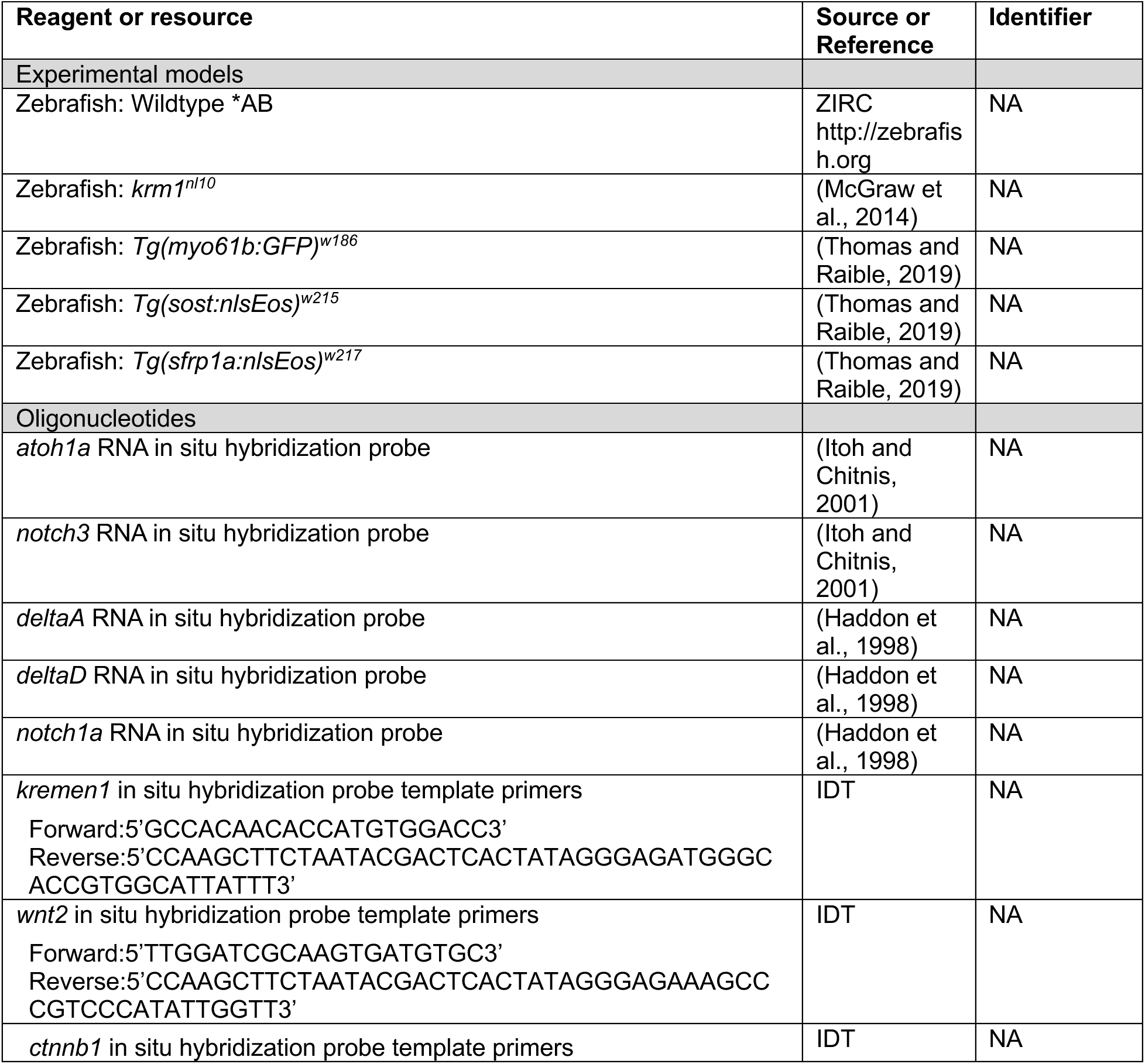
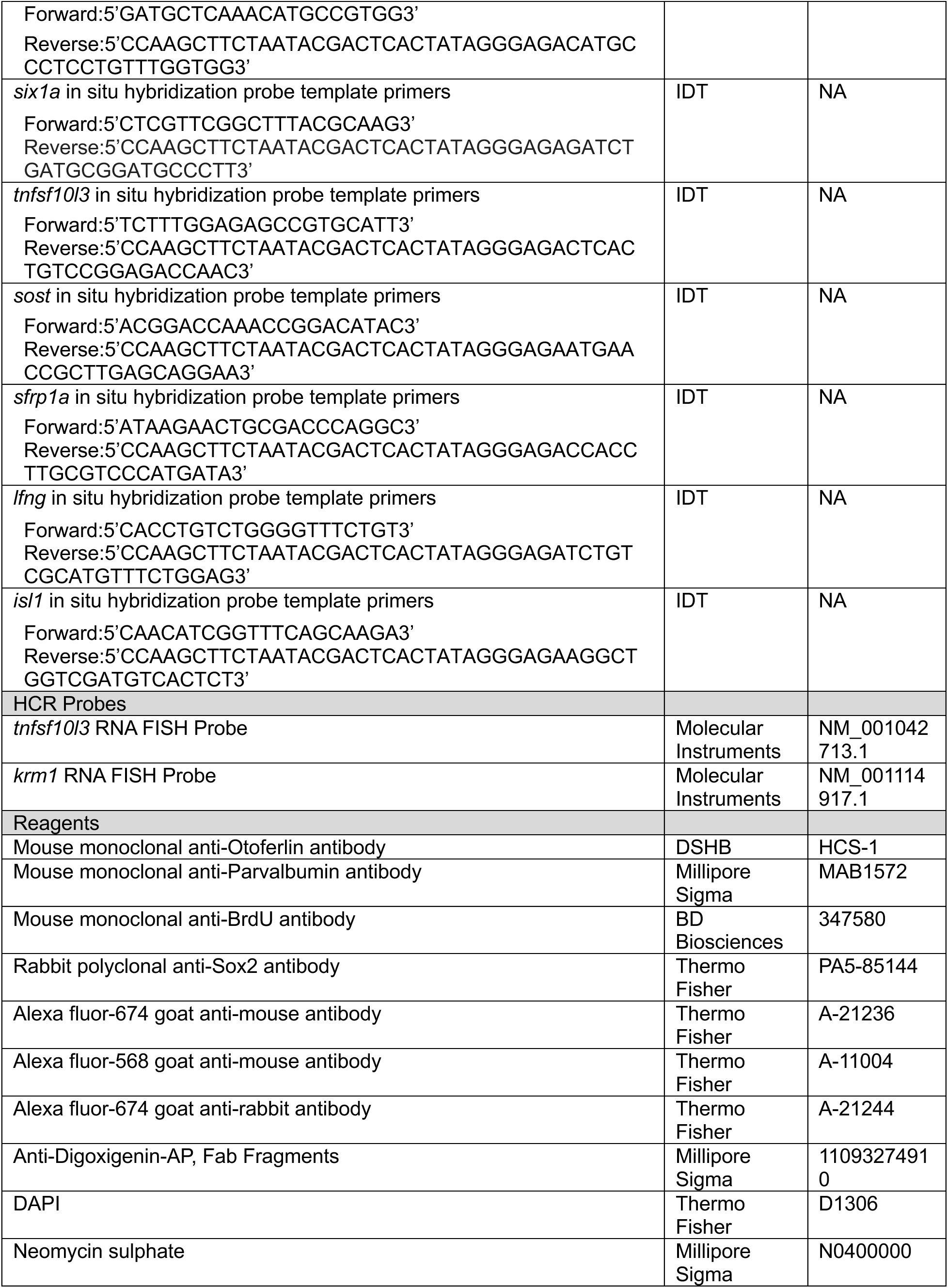
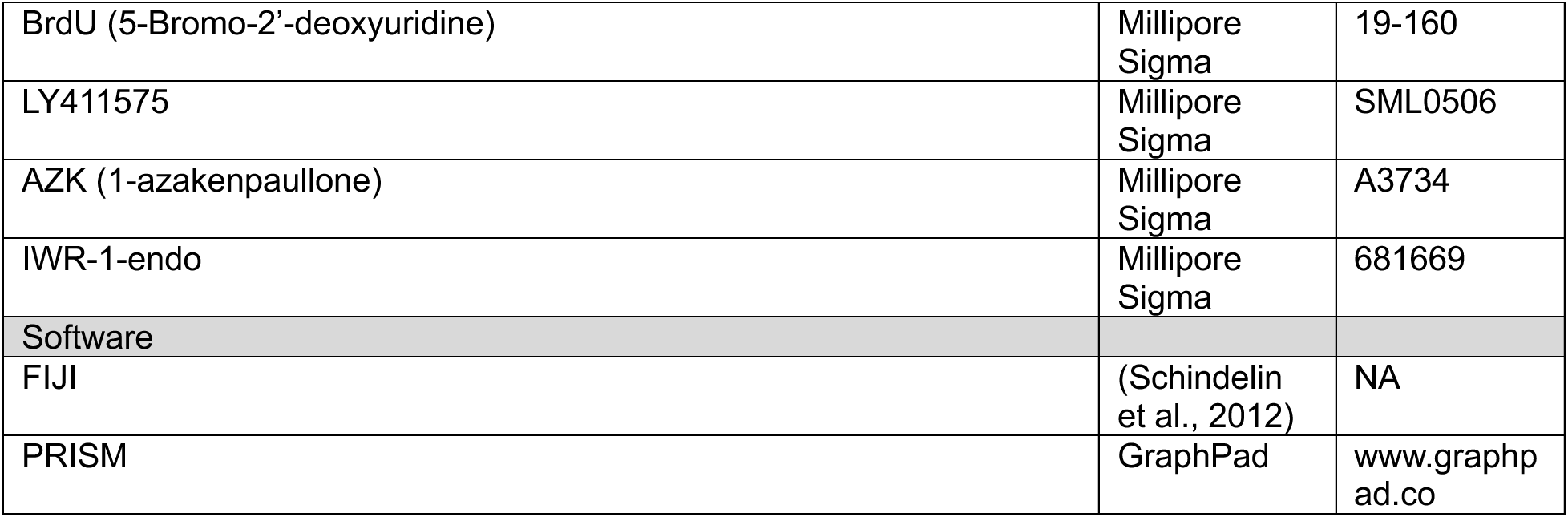
Key Resource Table.

### Zebrafish lines and maintenance

The following zebrafish lines were used: wild-type*AB (ZIRC;http://zebrafish.org), *Tg(myosin6b:GFP)^w186^* (^19^),*Tg(sost:nlsEos)^w215^* (^19^), *Tg(sfpr1a:nlsEos)^w217^* (^19^), and *krm1^nl10^* (^31^). Zebrafish were maintained and staged according to standard protocols (^48^). Larvae were kept in E3 embryo medium (14.97 mM NaCl, 500 μM KCL, 42 μM Na_2_HPO_4_, 150 μM KH_2_PO_4_, 1 mM CaCl_2_ dihydrate, 1 mM MgSO_4_, 0.714 mM NaHCO_3_, pH 7.2). For all experiments, larvae were treated with tricaine (Syndel) prior to fixation in 4% paraformaldehyde/PBS (Thermo Fisher). All work was performed in accordance with the McGraw laboratory protocol #41153 approved by the UMKC IACUC committee.

### RNA in situ hybridization and HCR fluorescent in situ hybridization

Whole mount RNA in situ hybridization was carried out using established protocols (^33^), modified with a 5-minute Proteinase K (Thermo Fisher) treatment to preserve neuromast integrity. The probes used were: *atoh1a*(^49^), *notch3*(^49^), *delta*(^50^)*, deltaD*(^50^)*, notch1a, kremen1*, *wnt2*, *ctnnb1*, *six1a*, *tnfsf10l3*, *sost, sfrp1a, lfng, and isl1.* Antisense probes were generated using established protocols (^33^) or using a PCR-based protocol (^51^). Hybridization chain reaction fluorescent in situ hybridization (HCR -FISH) was carried out following the manufacturer’s protocol. (Molecular Instruments). The probes used were *krm1*-B1(4pmol) and *tnfsf10l3-*B2 (4pmol), with the amplifiers B1-546 and B2-647 respectively (Molecular Instruments). Larvae were subsequently labeled with DAPI and mounted using Fluorescent Mounting Media (EMD Millipore) to prevent fading.

### immunohistochemistry and FM1-43FX labeling

Whole mount immunolabeling was performed using established protocols (Ungos et al., 2003). The primary antibodies used were: anti-Otoferlin antibody (mouse monoclonal, 1:200, DSHB, University of Iowa), anti-Sox 2 antibody (rabbit polyclonal, 1:100, Invitrogen) and anti-BrdU antibody (mouse monoclonal, 1:100, BD Biosciences). Secondary antibodies used were: goat anti-rabbit Alexa-647 (1:1000, Invitrogen), goat anti-mouse Alexa-568 (1:1000, Invitrogen), and goat anti-mouse Alexa-647 (1:1000, Invitrogen). Mature hair cells were labeled by a 1-minute incubation in 3μM FM1-43FX (Invitrogen; ^52^). Nuclei were labeled with 30μM DAPI (Thermo Fisher).

### Regeneration and BrdU incorporation

For hair cell ablation, 5 days post fertilization (dpf) larvae were incubated in 400 μM neomycin (NEO, Millipore-Sigma) for 0.5 hours and then washed 3x in fresh embryo medium. In experiments analyzing complete regeneration, larvae were collected 3 days following NEO exposure. For RNA in situ hybridization, larvae were collected 4 hours or 1 day post NEO. To assess multiples rounds of hair cell ablation and regeneration, larvae were exposed to NEO at 5 dpf, 8 dpf, 11 dpf, and 14 dpf; larvae were fixed 3 days after their final NEO exposure. Cellular proliferation during regeneration was analyzed using bromodeoxyuridine (BrdU; Millipore Sigma). BrdU incorporation was carried out using established protocols (^53, 54^); larvae were incubated in 10 mM BrdU for 24 hours at set times following NEO-induced ablation: for 5-6 day BrdU, larvae were exposed to BrdU immediately after NEO for 24 and then transferred to fresh E3 for 2 days; for 6-7 day BrdU, larvae were kept in E3 for 1 day following NEO, then incubated in BrdU for 1 and transferred to E3 for 1 day; and for 7-8 day BrdU, larvae were kept in E3 for 2 days following NEO exposure, followed by by 24 hours BrdU incorporation. All larvae were collected for fixation at 8 pdf.

### Inhibitor treatment conditions

Wnt signaling was activated using the Gsk3β inhibitor 1-azakenpaullone (Azk; Millipore Sigma; ^38^) or was inhibited using the Tankyrase inhibitor IWR-1-endo (Millipore Sigma), both were dissolved in DMSO and diluted in E3 embryo medium to 9μM and 22μM respectively. Control larvae were incubated in less than 1% DMSO. Notch inhibition was done using the ψ-secretase inhibitor LY411575 (Millipore Sigma) at 50μM with DMSO as a control. For regeneration experiments in the presence of inhibitors, 5 dpf larvae were treated with inhibitors or DMSO for 5 hours prior to exposure to NEO. Following NEO-induced ablation, larvae were incubated in DMSO or inhibitors for 3 days of regeneration.

### Photoconversion

For photoconversion experiments using *Tg(sost:nlsEos)^w215^* or *Tg(sfrp1a:nlsEos)^w217^* fish, 5 dpf larvae were placed in a shallow depression slide and exposed to 405 nm light for 20 seconds using a Zeiss Imager.D2 compound microscope and a 10x objective. For regeneration experiments, photoconversion was carried out prior to NEO exposure. For live imaging of *Tg(sost:nlsEos)^w215^* fish, larvae were anesthetized using tricaine and embedded in 1.2% low melt agarose/E3 embryo medium.

### Image collection and data analysis

For imaging of RNA in situ hybridization and immunohistochemistry, processed larvae were placed in 50% glycerol/PBS and mounted on slides. For imaging of HCR in situ hybridization, larvae were mounted on slides in Fluorescent Mounting Media (Calbiochem) and imaged within 3 days of processing to prevent signal loss. Images were collected using a Zeiss 510 meta confocal microscope using Zen 2009 software. Images were processed using Fiji software (^55^) and brightness and contrast were adjusted using Photoshop (Adobe). Neuromasts L1-L4 were analyzed for each fish and cells were manually counted under blinded conditions. All statistical analysis was carried out using Graphpad Prism 10 (GraphPad Prism version 10.0.0 for Mac, GraphPad Software, San Diego, California USA, www.graphpad.com). A Student’s *t-* test was used for pair-wise comparisons where samples passed a normality test and a One-way ANOVA with Dunn’s multiple comparisons was used when there were more than 2 conditions. Significance was set at p<0.05. All data is presented as ± standard deviation (SD).

## Author Contributions

Conceived and designed experiments: EM, MK, JB, HFM. Preformed experiments: EM, MK, BL, JB, HFM. Analyzed date: EM, MK, JB, HFM. Wrote Manuscript: HFM. Edited Manuscript: JB, HFM.

## Acknowledgements

The authors thank Dr. David Raible of the *Tg(sost:nlsEos)^w215^* and *Tg(sfrp1a:nlsEos)^w217^* zebrafish lines. This work was supported by funding provided to HFM by NIGMS (R16GM146690; https://www.nigms.nih.gov/).

